# The role of Dicer-dependent RNA interference in regulating cross-species communication during fungus-fungus interactions

**DOI:** 10.1101/2021.06.28.450161

**Authors:** Edoardo Piombo, Ramesh Raju Vetukuri, Anders Broberg, Pruthvi B. Kalyandurg, Sandeep Kushwaha, Dan Funck Jensen, Magnus Karlsson, Mukesh Dubey

## Abstract

Dicer-like (DCL) proteins play a vital role in transcriptional and post-transcriptional gene silencing, also known as RNA interference (RNAi), by cleaving double-stranded RNAs or single-stranded RNAs with stem-loop structures into small RNAs. Although DCL-mediated RNAi can regulate interspecific communication between pathogenic/mutualistic organisms and their hosts, its role in parasitic fungus-fungus interactions is yet to be investigated. In this study, we deleted *dcl* genes in the mycoparasitic fungus *Clonostachys rosea* and analyzed the transcriptome and secondary metabolome to characterize the regulatory functions of DCL-dependent RNAi in mycoparasitism. Deletion of *dcl2* resulted in a mutant with reduced growth rate, pigment production and antagonism towards the plant pathogenic fungus *Botrytis cinerea*. Moreover, the Δ*dcl2* mutant displayed a reduced ability to control fusarium foot rot disease on wheat, caused by *Fusarium graminearum*, and reduced production of 62 secondary metabolites (SM) including yellow-coloured sorbicillinoids. Transcriptome sequencing of the *in vitro* interaction between the *C. rosea* Δ*dcl2* strain and *B. cinerea* or *F. graminearum* identified downregulation of genes coding for transcription factors, membrane transporters, hydrolytic enzymes and SM biosynthesis enzymes putatively involved in antagonistic interactions, in comparison with the *C. rosea* wild type interaction. Sixty-one putative novel microRNA-like RNAs (milRNAs) were identified in *C. rosea*, and 11 was upregulated in the Δ*dcl2* mutant. In addition to putative endogenous gene targets, these DCL2-dependent milRNAs were predicted to target *B*. *cinerea* and *F. graminearum* virulence factor genes, which showed an increased expression during interaction with the Δ*dcl2* mutant incapable of producing the targeting milRNAs. This paper constitutes the first step in elucidating the role of RNAi in mycoparasitism, with important implications for biological control of plant diseases. This study further indicates a possible cross-species regulatory activity of fungal milRNAs, emphasizing a novel role of RNAi in fungal interactions and ecology.

**Author summary:** RNA interference (RNAi) is a conserved cellular mechanism mediated by small RNAs (sRNAs) regulating biological processes through the targeted destruction or modulation of RNA filaments necessary for protein synthesis. Dicer-like endoribonucleases (DCL) play a vital role in the RNAi pathway by generating sRNAs. In this study, we identified two DCL-encoding genes in the mycoparasitic fungus *Clonostachys rosea* and investigated a role of DCL-mediated RNAi in interference interactions between *Clonostachys rosea* and the two important fungal pathogens *Botrytis cinerea* and *Fusarium graminearum* (here called mycohost). Using transcriptome (sRNA and mRNA) sequencing and secondary metabolome analysis approach, we found that the *dcl* mutants were not able to produce 11 sRNAs predicted to finetune the regulatory network of genes known to be involved in production of hydrolytic enzymes, antifungal compounds, and membrane transporters needed for antagonistic action of C. *rosea*. We also found *C*. *rosea* sRNAs putatively targeting known virulence factors in the mycohost, indicating RNAi-mediated cross-species communication. Our study expanded the understanding of underlying mechanisms of cross-species communication during interference interactions and showed that DCL-mediated RNAi is an important regulator of parasitic fungus-fungus interactions. The results pose the base for future works studying the role of DCL-based cross-species RNAi in fungal interactions.

## 1. Introduction

Small RNAs (sRNAs) are a group of non-coding RNA. They play a central role in gene silencing at the transcriptional level through chromatin modification, and at the post-transcriptional level through targeted destruction of mRNAs, also known as RNA interference (RNAi) [1–5]. The central enzymatic components of RNAi pathways are dicer or dicer-like endoribonuclease (DCL), argonaute (AGO) protein (an sRNA-guided endonuclease) and RNA-dependent RNA polymerase (RDRP). DCL cleaves the double-stranded RNA precursors and single-stranded RNA precursors with hairpin structures to generate sRNAs, often ranging in size from 18 to 40 nucleotides, called small-interfering RNAs (siRNAs) and microRNAs (miRNAs; microRNA-like RNAs [milRNAs] in fungi), respectively. The resulting sRNA duplexes are then loaded in the AGO protein of the RNA-induced silencing complex (RISC). After removing the passenger strand of the sRNA duplex, the single strand sRNA within the AGO protein guides the RISC complex to the target and triggers silencing of genes, with fully or partially complementary sequences, by inhibiting translation or inducing transcript cleavage of the sRNA/target duplex [1–5]. An RDRP, that is used to generate dsRNAs from aberrant RNAs, amplifies the silencing signal through the formation of secondary siRNAs. In fungi, the most studied RNAi pathways are mediated by siRNAs and milRNAs, and are dependent on DCL for biogenesis and are thus called dicer-dependent RNAi. Dicer-independent RNAi such as those mediated by dicer-independent small interfering RNAs (disiRNAs) has also been identified in the filamentous fungus *Neurospora crassa* [6].

Small-RNA mediated RNAi is an evolutionarily conserved process of self-defense triggered by a wide variety of exogenous nucleic acids such as invading viruses, transgenes, transposons, and plasmids [7, 8]. In fungi, a role of sRNA-mediated RNAi pathways in genome defence against the insertion of repetitive transgenes during vegetative growth (quelling) and sexual phase of the life cycle [meiotic silencing of unpaired DNA (MSUD)] was first reported in *N*. *crassa* [9–11]. Since then, RNAi pathways and their role in genome defence against retrotransposon activity have been demonstrated in several fungal species with diverse lifestyles [8,12–20]. However, in some fungal species such as *Saccharomyces cerevisiae* and *Ustilago maydis*, genes related with the RNAi pathways are absent [21, 22]. In addition to the role of genome defence against transgenes, the fungal RNAi machinery generates a variety of sRNAs that are involved in the regulation of numerous biological processes through targeted gene silencing [8, 23]. For instance, sRNAs (mainly milRNAs) are found to be differentially expressed in fungi during different growth phases, developmental stages, and environmental conditions including those involved in host-pathogen interactions [24–34]. Furthermore, sRNAs can move bidirectionally between the species and modulate cellular functions of recipient cells by hijacking their RNAi machinery. Thus they play an important role in interspecies communication between closely interacting symbiotic organisms, including parasitic and mutualistic interactions [35–40]. However, the role of sRNA in parasitic fungus-fungus interactions is yet to be investigated.

The filamentous fungus *Clonostachys rosea* is a ubiquitous soil-borne ascomycete with a complex lifestyle as a necrotrophic mycoparasite by killing other fungi and feeding on dead mycelia, as a saprotroph by digesting organic material in soil and as a beneficial plant root coloniser [41]. *C. rosea* efficiently overgrows and kills its mycohosts such as *Botrytis cinerea* and *Fusarium graminearum* [41–43]. During mycoparasitic interactions or exposure to the secreted factors from mycohosts, *C. rosea* induces expression of genes associated with the production of secondary metabolites, hydrolytic enzymes and other secreted proteins [43–50]. Furthermore, *C*. *rosea* induces expression of genes coding for membrane transporters to efflux the endogenous toxic compounds and exogenous metabolites that may come from interacting organisms during the interspecific interactions [49,51,52]. The role of secreted proteins/enzymes, secondary metabolites, and membrane transporters in antibiosis and mycoparasitism in *C. rosea* is proven [42–44,50,53,54], however, the role of RNAi in regulating the cellular regulatory network during such interactions is yet to be investigated.

The present work aims to i) characterize the RNAi machinery in *C*. *rosea*; ii) identify milRNAs that are key regulators of genes associated with the antagonistic/mycoparasitic activity in *C*. *rosea*, and their potential endogenous and cross-species gene targets; and iii) investigate common or species-specific responses in sRNA-mediated gene regulation in *C*. *rosea* against mycohosts. We used the two important plant pathogenic fungi *B*. *cinerea* and *F*. *graminearum* as different mycohosts, as they are taxonomically different from each other and represent different disease types on different crops. We hypothesized that i) sRNAs regulate mycoparasitic interactions in *C*. *rosea* at endogenous and cross-species level; ii) *C*. *rosea* responds with both common and mycohost-specific reactions towards *B*. *cinerea* and *F*. *graminearum*. To test the hypotheses, we generated gene deletion and complementation strains of genes coding for DCL proteins (DCL1 and DCL2) in *C*. *rosea* and used a holistic approach (sRNA, transcriptome and secondary metabolome analysis) to investigate the sRNA-mediated regulatory network and its influence on mycoparasitic fungus-fungus interactions at endogenous and cross-species level.

## 2. Material and methods

### 2.1 Fungal strains and culture conditions

*C. rosea* strain IK726 wild type (WT) and mutants derived from it, *B. cinerea* strain B05.10 and *F. graminearum* strain PH1 were used in the study. The fungal cultures were maintained on potato dextrose agar (PDA; Oxoid, Cambridge, UK) medium at 25°C.

### 2.2 Gene identification and phylogenetic analysis

The *C*. *rosea* strain IK726 genome version 1 [41] and version 2 [55], were screened for the presence of genes encoding DCL, AGO and RDRP by BLASTP analysis. Presence of conserved domains were analysed with Simple Modular Architecture Research Tool (SMART) [56], InterProScan [57] and conserved domain search (CDS) [58].

Amino acid sequences of DCLs (DCL1 and DCL2), AGOs (AGO1 and AGO2) and RDRPs of several fungal species (Table S1), were retrieved from the Uniprot and Genbank databases [59, 60]. The sequences of Dicer1, Argonaute 1 and RDRP1 of the model plant *Arabidopsis thaliana* were retrieved from the Uniprot database [59] and used as outgroups. Sequences were aligned with mafft v.7 [61] with options suggested for less than 200 sequences (L-INS-i), and the phylogenetic trees were generated with iqtree v.1.6.12 [62] with 1000 bootstrap replicates and option “MFP” to find the best substitution model. Figtree v.1.4.4 [63] was used to visualize the trees.

### 2.3 Construction of deletion vector, transformation, and mutant validation

The ∼ 1 kb 5’-flank and 3’-flank regions of *dcl1* and *dcl2* were amplified from genomic DNA of *C*. *rosea* using gene specific primer pairs (Table S2) as indicated in Figure S1 [53]. Gateway cloning system (Invitrogen, Carlsbad, CA) was used to generate entry clones of the purified 5’-flank and 3’-flank PCR fragments as described by the manufacturer (Invitrogen, Carlsbad, CA). The hygromycin resistance cassette (hygB) generated during our previous studies [43, 64] from pCT74 vector, as well as a geneticin resistance cassette generated as a PCR product from the pUG6 vector [65], were used. Three-fragments multisite gateway LR recombination reaction was performed using entry plasmid of respective fragments and destination vector pPm43GW [66] to generate the deletion vectors. Complementation cassettes for *dcl1* and *dcl2* were constructed by PCR amplification of the full-length sequence of *dcl1* and *dcl2* including around 800 bp upstream and around 500 bp downstream regions from genomic DNA of *C. rosea* WT using gene specific primers (Table S2). The amplified DNA fragments were purified and integrated into destination vector pPm43GW using two fragment gateway cloning technology to generate complementation vectors.

*Agrobacterium tumefaciens*-mediated transformation (ATMT) was performed based on a previous protocol for *C. rosea* [43, 67]. Transformed strains were selected on plates containing either hygromycin for gene deletion or geneticin for complementation. Validation of homologous integration of the deletion cassettes in putative transformants were performed using a PCR screening approach with primer combinations targeting the hygB cassette and sequences flanking the deletion cassettes (Figure S1), as described previously [45, 68]. The PCR positive transformants were tested for mitotic stability and were then purified by two rounds of single spore isolation [64]. To determine transcription of *dcl1* and *dcl2* in the WT, deletion and complementation strains, total RNA from the respective strains were isolated (Qiagen, Hilden, Germany). After DNase I treatment, according to the manufacturer’s instructions (Merck, Kenilworth, NJ) reverse transcription (RT) PCR was performed using RevertAid premium reverse transcriptase (Fermentas, St. Leon-Rot, Germany) and gene specific primer pairs (Table S2).

### 2.4 Phenotypic analyses

Phenotypic analyses experiments were performed with *C*. *rosea* WT, deletion strains *dcl1* (Δ*dcl1*) and *dcl2* (Δ*dcl2*), and their respective complemented strains Δ*dcl1*+ and Δ*dcl2*+. Each experiment was repeated two times with similar results.

The growth rate, colony morphology and conidia production were analysed in four biological replicates following procedures described before [43]. To analyse mycelial biomass, an agar plug of *C*. *rosea* strains were inoculated in 50 ml conical flasks with 20 ml PDB (Oxoid, Cambridge, UK) and incubated at 25 °C under constant shaking condition (100 rpm). Biomass production was determined by measuring mycelial dry weight 5 days post inoculation. Antagonistic behaviour against *B*. *cinerea* and *F. graminearum* was tested using an *in vitro* plate confrontation assay on PDA medium as described before [51]. The growth of *B*. *cinerea* and *F. graminearum* was measured daily until their mycelial front touched the *C*. *rosea* mycelial front. The experiments were performed in four biological replicates. The biocontrol ability of *C. rosea* strains against *F*. *graminearum* was evaluated in a fusarium foot rot assay as described previously [69]. In brief, surface sterilised wheat seeds were treated with *C. rosea* conidia (1e+07 conidia/ml) in sterile water, sown in moistened sand, and kept in a growth chamber after pathogen inoculation [51]. Plants were harvested three weeks post inoculation and disease symptoms were scored on 0-4 scale as described before [51, 69]. The experiment was performed in five biological replicates with 15 plants in each replicate.

### 2.5 Statistical analysis

Analysis of variance (ANOVA) was performed on phenotype data using a General Linear Model approach implemented in Statistica version 16 (TIBCO Software Inc., Palo Alto, CA). Pairwise comparisons were made using the Tukey-Kramer method at the 95% significance level.

### 2.6 Metabolite analysis

An agar plug of *C*. *rosea* strains was inoculated on PDA (Oxoid, Cambridge, UK) and allowed to grow for 10 days at 25°C. Agar plugs together with mycelia were harvested from the centre of plates using 50 ml falcon tubes [53]. The mycelial plug was sonicated for 15 min in 20 ml of methanol and then one ml of extract was transferred to a 1.5 ml centrifuge tube for centrifugation at 10000 × g for 5 min. Supernatants were collected, and analyzed by UHPLC-MS (Ultra-high performance liquid chromatography-mass spectrometry) on a reversed phase column (2.1 × 50 mm, 1.5 μm, Accucore Vanquish, Thermo Scientific, Waltham, MA) using a gradient of acetonitrile (MeCN) in water, both with 0.2% formic acid (10-95% MeCN in 3 min, 95% MeCN for 1.2 min, at 0.9 mL min^-1^). The MS was operated in positive mode with scanning of *m/z* 50-1500, and the mass spectra were calibrated against sodium formate clusters using the software Compass DataAnalysis 4.3 (Bruker Daltonics, Bremen, Germany) that was also used for general data analysis. UHPLC-MSMS was run on the same instrument, column and UHPLC-conditions, using the auto MSMS function (1+ precursor ions, *m/z* 50-1500, ramped fragmentation energies: 20/30/35 eV for *m/z* 200/500/1000). The UHPLC-MS data was converted to mzXML format using DataAnalysis 4.3 and ion-chromatogram peak picking in the range 5 s to 200 s was done by the program XCMS in the software environment R using the centWave method (peakwidth 3-20 s, *m/z* tolerance 5 ppm, noise 1000) [70, 71]. XCMS was used for subsequent peak grouping and missing peak filling. For each sample, the resulting molecular feature peak-areas were normalized against the sum of peak-areas, and the resulting relative peak-areas were 10-logarithmized. The data were used for principal component analysis (PCA) and ANOVA was used to evaluate significant differences in concentrations between strains. Tentative compound identification was done by comparison of high-resolution mass spectrometry (HRMS)-data on fungal compounds from the databases Antibase and combined chemical dictionary. The identity of the tentatively identified compounds was further corroborated by analysis of MSMS-data. The experiment was performed in five biological replicates.

### 2.7 Dual culture interaction experiment for sRNA and transcriptome sequencing

An agar plug of *C*. *rosea* strains was inoculated at edge of a 9-cm-diameter potato dextrose agar (PDA; Merck, Kenilworth, NJ) Petri plate covered with a durapore membrane filter (Merck, Kenilworth, NJ) for easy harvest of mycelia. The mycohost fungi *B. cinerea* or *F*. *graminearum* were inoculated at opposite side of the plate [43]. Due to different mycelial growth rates, *C*. *rosea* was inoculated 7 days prior to the inoculation of *F*. *graminearum* or *B*. *cinerea*. The mycelial front (5 mm) of *C*. *rosea* was harvested together with the mycelial front (5 mm) of *B*. *cinerea* (Cr-Bc) or *F*. *graminearum* (Cr-Fg) at hyphal contact stage of interactions (Figure S2) and snap frozen in liquid nitrogen. The experiment was performed in three biological replicates.

### 2.8 RNA extraction, library preparation and sequencing

Total RNA was extracted using the mirVana miRNA isolation kit following the manufacturer’s protocol (Invitrogen, Waltham, MA). The RNA quality was analyzed using a 2100 Bioanalyzer Instrument (Agilent Technologies, Santa Clara, CA) and concentration was measured using a Qubit fluorometer (Life Technologies, Carlsbad, CA). For sRNA and mRNA sequencing, the total RNA was sent for library preparation and paired-end sequencing at the National Genomics Infrastructure (NGI) Stockholm, Sweden. The sRNA library was generated using TruSeq small RNA kit (Illumina, San Diego, CA), while the mRNA library was generated using TruSeq Stranded mRNA poly-A selection kit (Illumina, San Diego, CA). The sRNA and mRNA libraries were sequenced on one NovaSeq SP flowcell with a 2×50 bp reads and NovaSeqXp workflow in S4 mode flow cell with 2×151 setup, respectively, using Illumina NovaSeq6000 equipment at NGI Stockholm. The Bcl to FastQ conversion was performed using bcl2fastq_v2.19.1.403 from the CASAVA software suite [72]. The quality scale used was Sanger / phred33 / Illumina 1.8+. The raw data was submitted to ENA in the bioproject PRJEB43636.

#### 2.8.1 Functional annotation of genomes

The predicted proteomes of *C. rosea* strain IK726, *B. cinerea* strain B05.10 (ASM14353v4) and *F. graminearum* strain PH-1 (ASM24013v3) were annotated through BLAST2GO v.5.2.5 [73] and InterProScan v.5.46-81.0 [57] with default parameters to identify transcription factors. Secondary metabolite clusters were predicted through antiSMASH v.5.0 [74], while predicted enzymes involved in the metabolism of carbohydrates (CAZymes) were identified through the dbCAN2 meta-server [75]. The amino acid sequences of *B*. *cinerea* and *F*. *graminearum* were compared with the PHI-base database using BLAST [76] with minimum 80% in both identity and query coverage. All identified matches described in the PHI-base annotation by the keywords “reduced virulence” or “loss of pathogenicity” were considered to be potential virulence factors.

#### 2.8.2 Differential expression and GO enrichment analyses

Reads were trimmed with bbduk v.38.86 [77] with the following options:

bbduk.sh in1=read1.fastq in2=read2.fastq out1=read1_clean.fastq out2=read2_clean.fastq ref=reference.fa ktrim=r k=23 mink=11 hdist=1 tpe tbo qtrim=r trimq=10

Successful cleaning and adapter removal was verified with fastqc v. 0.11.9 (https://www.bioinformatics.babraham.ac.uk/projects/fastqc/). Since all the samples represented the interaction of two organisms, the genome of *C. rosea* was concatenated with the one of either *B. cinerea* or *F. graminearum*, creating two “combined genomes” (Cr-Fg and Cr-Bc), and the same was done with the annotations in gff format. Reads from *C. rosea*-*B. cinerea* interaction were aligned to the Cr-Bc genome, while reads from *C. rosea*-*F. graminearum* interaction were aligned to the Cr-Fg. The chosen aligner was STAR v.2.7.5c [78], with default options, and the count tables were then generated through featureCounts v.2.0.1 [79]. Finally, the differential expression analysis was done with DESeq2 v.1.28.1 [80], where a FDR < 0.05 in combination with a log2FoldChange > 1.5 or < −1.5 was considered to define differentially expressed genes (DEGs). Enrichment in gene ontology (GO) terms of DEGs was determined through Fisher tests integrated in the BLAST2GO suite, with an FDR threshold of 0.05.

#### 2.8.3 Mapping of sRNAs

sRNA reads were trimmed with bbduk v.38.86 [77] with the same options used for mRNA read trimming, and successful cleaning and adapter removal was verified with fastqc v. 0.11.9. The program SortMeRna v.4.2.0 [81] was used to remove structural sRNA (rRNA, tRNA, snRNA, and snoRNA) from the reads, and sequences within the 18-32 bp length range were isolated with the command reformat.sh of the BBTools suite [77]. The database of structural RNAs used for SortMeRna consisted in the rRNA sequences of the SILVA database [82], while snRNA, tRNA and snoRNA sequences were downloaded from the NRDR database [83]. After filtering, the sRNA reads were mapped to the Cr-Bc and Cr-Fg genomes with STAR, with options recommended for sRNA mapping:

STAR --genomeDir index/ --readFilesIn read1.fq read2.fq --outFileNamePrefix prefix --outFilterMismatchNoverLmax 0.05 --outFilterMatchNmin 16 --outFilterScoreMinOverLread 0 --outFilterMatchNminOverLread 0 --alignIntronMax 1 --alignEndsType EndToEnd As per STAR default option, reads with good mapping results on more than 20 different loci were considered as not mapped.

Untranslated regions (UTRs) and introns were added to the gff files of the genomes through “add_utrs_to_gff” (https://github.com/dpryan79/Answers/tree/master/bioinfoSE_3181) and GenomeTools with the “-addintrons” option [84], respectively. Promoters were added as well through an *ad hoc* python script (https://github.com/EdoardoPiombo/promoter_extractor), considering promoters to be composed of the first 1000 bases upstream of a gene, or of all the bases until the end of the precedent gene. Introns, promoters and UTRs were all considered when featureCounts was used to generate the count tables.

#### 2.8.4 Prediction of miRNA-like RNAs and following analyses

Putative milRNAs were predicted with mirdeep2 v.2.0.1.2 [85]. The miRbase database [86], as well as all the fungal milRNA sequences from RNAcentral [87], were used to provide reference sequences from other species. To ensure the novelty of newly detected milRNAs, BLAST was used to compare them with the fungal milRNAs identified in several other studies, plus all the fungal milRNAs available in RNAcentral, requiring 95% minimum identity and query coverage [25,34,87–91]. Non-structural sRNA reads, previously mapped to the genomes with STAR, were counted with featureCounts and the differentially expressed milRNAs were identified with DESeq2, with the same thresholds used for DEGs analysis.

The sRNA_toolbox [92] was used to predict putative targets for the identified milRNAs. The prediction was carried out with the animal-based tools PITA, Miranda, TargetSpy [93–95] and simple seed analysis, and with the plant-based tools psRobot, TAPIR FASTA and TAPIR RNAhybrid [96, 97]. Target-milRNA couples identified by at least 3 animal-based tools or 2 plant-based ones were retained for the following analyses. Predicted targets were retained only when they had a significant expression (FDR < 0.05) with Log2FC >1.0 opposite to the milRNA. Putative targets of downregulated milRNAs were therefore considered only when they were overexpressed. Afterwards, predicted targets present in double copy in their genome were removed from the analysis. Repetitive elements in the genome of *C. rosea* were then predicted following the guidelines for basic repeat library construction (http://weatherby.genetics.utah.edu/MAKER/wiki/index.php/Repeat_Library_Construction--Basic), using all fungal transposons in RepetDB as known transposons [98], and putative milRNA targets within 700 bp from any *C. rosea* transposon were removed from the analysis.

#### 2.8.5 Validation of milRNA-expression through stem-loop RT-qPCR

milRNA specific stem-loop RT-qPCR primers (Table S2) were designed as described previously [99]. One μM stem-loop RT primers were denaturated at 65°C for 5 min, and immediately transferred to ice. For each milRNA RT reaction, a “no RNA” master mix was prepared with 0.5 μl of 10 mM dNTP (Thermo Scientific, Waltham, MA), 5x SSIV buffer, 2 μl of 0.1 M DTT, 0.1 μl of RiboLock RNase inhibitor (40U/μl), 0.25μl of SSIII reverse transcriptase (Invitrogen, Waltham, MA), 1 μl of denaturated stem-loop RT primer, and 1 μl of 5 μM reverse primer of *C. rosea* actin (*act*) reference gene. Ten ng of RNA template used for NGS analysis was added into respective reactions. The tubes were then incubated in a thermal cycler at 16°C for 30 min, followed by 60 cycles of pulsed RT at 30°C for 30 s, 42°C for 30 s and 50°C for 1 s, and enzyme inactivation at 85°C for 5 min. Quantitative PCR was performed using DyNAmo Flash SYBR Green kit (Thermo Scientific, Waltham, MA) following manufacturer’s instructions. The Ct values of milRNA were normalized to that of *act* to be used for quantification using the ΔΔCT method [100].

## 3. Results

### 3.1 Identification and sequence analysis of the predicted RNAi machinery in *C*. *rosea*

Genes coding for different protein components involved in the RNAi pathway were identified through BLAST analysis of *C*. *rosea* strain IK726 genome ver. 1 [41] and ver. 2 [55] using *N. crassa* and *Trichoderma atroviride* AGO, DCL, and RDRP gene sequences as queries. Two AGO (AGO1, protein ID CRV2T00002735; AGO2, protein ID CRV2T00000975), two DCL (DCL1, protein ID CRV2T00009872; DCL2 protein ID CRV2T00008135) and three RDRP (RDRP1, protein ID CRV2T00001186; RDRP2, Protein ID CRV2T00002170; RDRP3, protein ID CRV2T00007201) genes were identified in the *C*. *rosea* genome. Analysis of the translated amino acid sequences for the presence of conserved modules identified the domains known to be present in DCL (DEXDc, HELICc, Dicer dimer, RNAse III), AGO (ArgoN, DUF, PAZ, ArgoL2, PIWI) and RDRP (RDRP) proteins (Figure S3). The characteristics of *C*. *rosea* AGOs, DCLs and RDRPs are presented in Table S3.

Phylogenetic analyses using DCL, AGO and RDRP amino acids sequences revealed that *C*. *rosea* putative DCLs were most closely related to their homologs in *Acremonium chrysogenum*, with around 57% sequence identity, and the same was true for *C. rosea* homologs of AGO1 and AGO2, but with an identity around 51%. The three putative RDRP genes were similar to their homologs in *A. chrysogenum* as well, with an identity of 37%, 42% and 55%, respectively. In the phylogenetic analyses, the putative DCLs of *C. rosea* diverged in two clusters separating the DCL1 and DCL2 from the analyzed species (Figure S4A), and the same was evident for AGO1 and AGO2 (Figure S4B). The tree generated from the RDRP sequences formed by three main clusters, each containing one of the *C. rosea* proteins (Figure S4C). Our data therefore suggests that *C*. *rosea* contain 2 DCL, 2 AGO and three RDRP genes, with clear orthologs in related species.

### 3.2 Generation of gene deletion and complementation strains

To investigate the biological roles of RNAi in *C*. *rosea*, genes encoding DCL proteins were selected for gene deletions as they act upstream in the RNAi pathways. Single *dcl1* and *dcl2* deletion strains (Δ*dcl1* and Δ*dcl2*) were generated and they were successfully complemented with *dcl1* and *dcl2*, respectively, to generate complementation strains Δ*dcl1*+ and Δ*dcl2*+. Results describing validation of gene deletion and complementation strains are presented in figure S1. Phenotypic analyses experiments were performed with *C*. *rosea* WT, *dcl* deletion strains (Δ*dcl1*, Δ*dcl2*) and their respective complemented strains Δ*dcl1*+ and Δ*dcl2*+.

### 3.3 Deletion of *dcl* affects growth, conidiation, antagonism and biocontrol

The growth rate of the Δ*dcl2* strain was 14% lower (*P* < 0.001) than the WT growth rate on PDA, while no significant difference was found between the Δ*dcl1* strain and the WT (Figure 1A). No significant difference in mycelial biomass (*P* ≤ 0.36) between the *C*. *rosea* WT and the *dcl* deletion strains was found (Figure S5A). We quantified the conidiation of *C*. *rosea* WT and deletion strains 24 days post inoculation (dpi). At this time, the colony perimeter of each strain had reached the edge of the 9 cm Petri dish. Conidia production of the Δ*dcl1* strain was 70% higher (*P* = 0.014) compared to that of the WT, while, no significant (*P* = 0.75) difference in conidia yield was recorded in the Δ*dcl2* strain (Figure 1B). Complementation strain Δ*dcl1*+ strains showed partial restoration of the conidial production phenotype observed in Δ*dcl1.* Morphological examination during growth on PDA revealed that the Δ*dcl2* had reduced ability to produce yellow pigment, while this phenotype remained unaffected in the Δ*dcl1* (Figure 1C). No other marked difference in colony morphology was observed between the WT and the *dcl* deletion strains.

**Figure 1:**
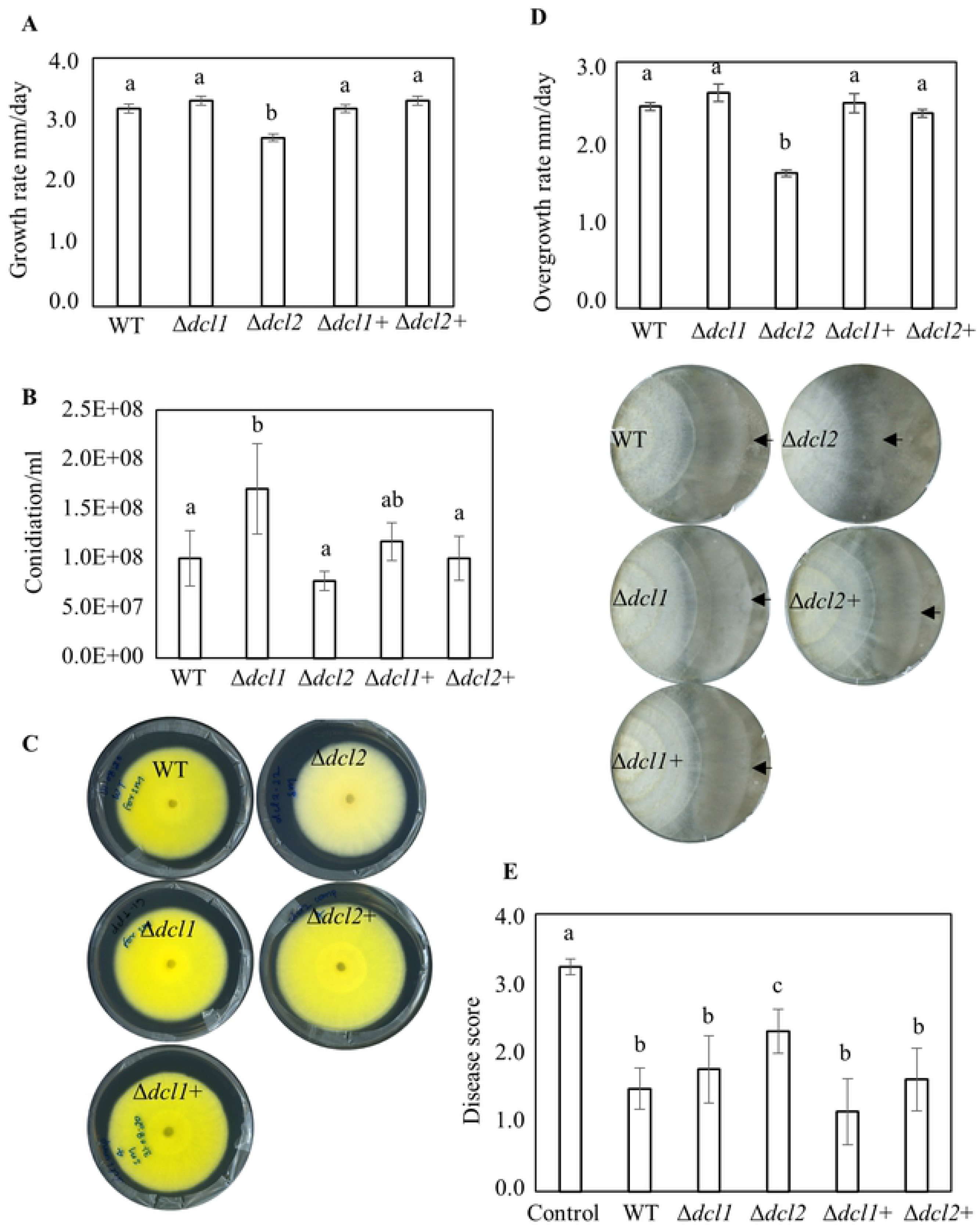
Phenotypic characterizations of *C*. *rosea* WT, deletion and complementation strains. **(A)** Growth rate of WT, *dcl* deletion and complemented strains. Strains were inoculated on PDA medium, incubated at 25°C and the growth rate was recorded 5 days post-inoculation (dpi). Error bars represent standard deviation based on 4 biological replicates. **(B)** Conidiation of WT, *dcl* deletion and complementation strains on PDA medium 24 dpi. Conidia were harvested in equal volume of water and were counted using a Bright-Line Haemocytometer as per instruction of manufacturer. Error bars represent standard deviation based on 4 biological replicates. **(C)** Deletion of *dcl2* affects pigment production in *C*. *rosea*. Strains were inoculated on PDA medium and were incubated at 25°C. The experiment of performed in four biological replicates and photographs of representative plates were taken 16 dpi. **(D)** Dual culture assay to test antagonistic ability of *C*. *rosea* WT, deletion and complementation strains against *B*. *cinerea*. Agar plugs of *C*. *rosea* strains (left side in the plate) and *B*. *cinerea* (right side in the plate) were inoculated on opposite sides in 9 cm diameter agar plates and incubated at 25°C. Growth rate (overgrowth) of *C*. *rosea* WT, deletion and complementation strains on *B*. cinerea was measured from the point of mycelial contact. The experiment was performed in four replicates and photographs of representative plates were taken 21 dpi of *C. rosea* strains. An arrowhead indicates the mycelial front of *C. rosea* strains. **(E)** *In vivo* assay to test the biocontrol ability of *C. rosea* strains against *F*. *graminearum* foot rot disease on wheat. Seeds were coated with *C. rosea* conidia and planted in moist sand together with a *F*. *graminearum* agar plug. Seedlings were harvested three weeks post inoculation and disease symptoms were scored on 0-4 scale. The experiment was performed in five biological replicates with 15 plants in each replicate. Different letters indicate statistically significant differences based on Tukey HSD method at the 95% significance level.

An in *vitro* dual culture assay was used to test whether deletion of *dcl1* or *dcl2* affected the antagonistic ability of *C. rosea*. No differences in growth rate of *F*. *graminearum* or *B*. *cinerea* were recorded during *in vitro* dual plate confrontation with either of the *dcl* deletion strains, compared with the WT (Figure S5B). However, a reduced ability (*P* < 0.001) to overgrow *B*. *cinerea* was observed in the Δ*dcl2* strains compared with the WT (Figure 1D). The growth rate of Δ*dcl2* strains displayed 33% reduction on *B*. *cinerea* mycelium (overgrowth rate) compared to the growth rate of WT (Figure 1D). In contrast, overgrowth of *F*. *graminearum* was not compromised in either of the deletion strains. However, a change in *F. graminearum* colour (pigment) was visible at the bottom side of the Δ*dcl2-F. graminearum* interaction zone (Figure S5C). In contrast to *in vitro* antagonism tests, a bioassay for biocontrol of fusarium foot rot diseases on wheat caused by *F*. *graminearum* displayed a significant 56% increase (*P =* 0.023) of disease severity in wheat seedlings previously seed coated with the Δ*dcl2* strain as compared with seedlings from seeds coated with *C. rosea* WT (Figure 1E). However, disease symptoms on seedlings from seeds coated with the Δ*dcl1* strains showed no significant difference compared with the WT.

### 3.4 Analysis of metabolites

The metabolites produced by the WT, *dcl* deletion and complementation strains were analysed by UHPLC-MS and -MSMS (Table S4). When analysing the UHPLC-MS data by PCA, the samples from the Δ*dcl1,* Δ*dcl1+* and WT strains grouped separated from each other (Figure 2A, left), and likewise the Δ*dcl2* and WT samples clustered separately (Figure 2A, right). The Δ*dcl2+* samples, however, clustered with the WT samples indicating restoration of metabolite production in Δ*dcl2+* strains. Two compounds were found to be present in significantly lower amount in the Δ*dcl1* and their production was restored in the Δ*dcl1*+ strains, along with 15 further compounds (ANOVA, FDR ≤ 0.01, Figure S6A, Table S4). Fifty-four metabolites were found to be present in significantly lower amounts in the Δ*dcl2* strain compared to the WT, and at the same time their production was restored in the Δ*dcl2+* strain (ANOVA, FDR ≤ 0.01, Figure S6B, Table S4). Seventeen of these compounds were tentatively identified or assigned to a compound class by UHPLC-MS, -MSMS and database mining (Figure 2B, Figure S7). A majority of these substances were monomeric or dimeric hexaketides of sorbicillin-type (e.g. sorbicillin, sorbicillinol, oxosorbicillinol, epoxysorbicillinol and bisvertinolone), whereas three glisoprenins (A, C and D) also were identified. The identification of some of these compounds is outlined below.

**Figure 2:**
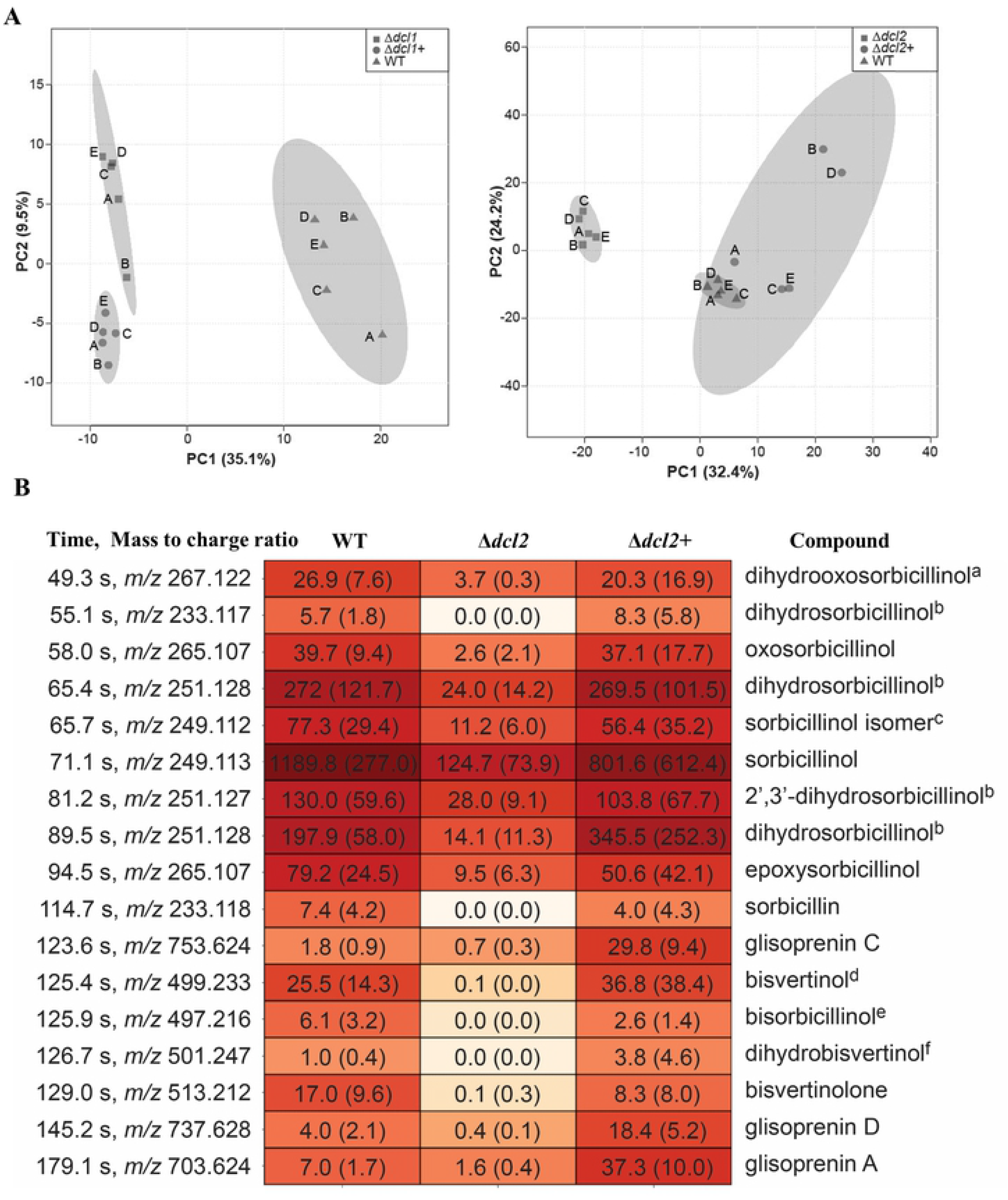
UHPLC-MS analysis of cultures of *C. rosea* WT and deletion strains. **(A)** Principal component analysis of UHPLC-MS data from analysis of metabolites produced by *C. rosea* WT and mutant strains (Δ*dcl1,* Δ*dcl2,* Δ*dcl1+* and Δ*dcl2+*). Shaded areas indicate 95% confidence regions. **(B)** Retention times, mass-to-charge ratios (*m/z*), extracted-ion chromatogram peak areas and tentative identification by UHPLC-MS and -MSMS of 17 metabolites produced in significantly lower amount in Δ*dcl2* mutants compared to the WT, and restored in Δ*dcl2+* compared (ANOVA FDR <0.01). The compound at 125.4 s was comparably underproduced-and restored also in the *Δdcl1* strains. Ions are [M+H]^+^ except for the compound at 55.1 s, which is [M+H-H_2_O]^+^. Peak areas shown are average peak areas × 10^-3^ with standard deviations in brackets. The heatmap is based on sum-normalized and 10-logaritmized peak areas. a) May also be dihydroepoxysorbicillinol. b) Proposed to be four different isomers of dihydrosorbicillinol. c) Has the same *m/z* as sorbicillinol, but different MSMS-data. d) May also be bisvertinoquinol or isobisvertinol. e) May also be bislongiquinolide or bisorbicillinolide or trichodimerol or trichotetronine. f) May also be isodihydrobisvertinol.

Sorbicillin was tentatively identified as a compound eluting at 114.7 s with [M+H]^+^ *m/z* 233.118, with two major fragment ions, *m/z* 95.049 and *m/z* 165.054, corresponding to bond cleavage on either side of the side-chain carbonyl (Figure S7). The ion at *m/z* 95.049 was found to be diagnostic for all monomeric and dimeric sorbicillin-type compounds containing a hexa-2,4-diene-1-one motif. Fragment ions corresponding to the ion with *m/z* 165.054 above, was important for all monomeric sorbicillin type compounds, and related fragment ions were frequently found with additional loss of CO and/or water, depending on the respective compound structure. The compound eluting at 71.1 s, with [M+H]^+^ *m/z* 249.113, was tentatively identified as sorbicillinol based on such fragment ions (Figure S7), and the two compounds at 58.0 s and 94.5 s, both with [M+H]^+^ *m/z* 265.207, were suggested to be oxosorbicillinol and epoxysorbicillinol, respectively, based on differences in fragment ions (Figure S7). Five compounds in figure 2B gave *m/z* values which after database mining suggested them to be vertinolide or hydroxyvertinolide, which are hexaketides similar to the sorbicillins, but with a lactone head-group instead of the aromatic ring or unsaturated cyclohexanone of sorbicillin-type compounds. In MSMS, however, the vertinolide-type compounds did not yield fragment ions supporting their structures. Instead, MSMS data suggested these compounds to be novel dihydrosorbicillinols or oxo/epoxy-dihydrosorbicillinol, respectively.

A large number of dimeric compounds of the sorbicillin-type is known [101] and several share the same molecular formula. These substances are dimerized by several different biosynthetic mechanisms, including Diels-Alder cycloaddition, Michael-type addition reactions and formation of hemi-ketals. The compound eluting at 129.0 s, with [M+H]^+^ *m/z* 513.212 (in accordance with the compound bisvertinolone) gave two major fragment ions at *m/z* 249.111 and *m/z* 265.107, both [M+H]^+^, corresponding to the constituting monomeric compounds of bisvertinolone, i.e. sorbicillinol and oxosorbicillinol, respectively (Figure S7). This pattern was observed for all putative dimeric sorbicillin-type compounds, i.e. in UHPLC-MSMS these compounds fragmented to yield ions of the presumed constituting monomeric compounds, and related ions after loss of CO and/or water (Figure S7). The formation of these fragment ions is possible for dimeric compounds formed by many different mechanisms, and therefore it was difficult to identify these by MSMS without access to authentic reference compounds or very detailed information about the MSMS behaviour of these compounds. Therefore, several alternative identities are listed in Figure 2B for some of the dimeric compounds. The polyhydroxy terpenes glisoprenin A, C and D were identified based on the *m/z* of their respective [M+H]^+^ ions, supported by the *m/z* of fragment ions (loss of multiple water molecules) detected in UHPLC-MSMS.

### 3.5 Transcriptome analysis of *Clonostachys rosea* WT and *dcl* deletion strains

To gain insights into the molecular mechanisms associated with the altered phenotypes of *C*. *rosea dcl* deletion strains, transcriptomes of *C*. *rosea* WT, Δ*dcl1* and Δ*dcl2* were analyzed by RNA-seq during the interactions with *B*. *cinerea* and *F*. *graminearum*. An average 20.5 million clean reads were obtained for each treatment. Since the sequences contained read pairs from both the interacting species, the reads originating from *C*. *rosea* or interacting mycohosts were identified by mapping to *C*. *rosea*, *B*. *cinerea* or *F*. *graminearum* genomes. During the *C. rosea*-*B. cinerea* interaction, on average 24% of reads were mapped to *C. rosea* genes, while 58% of reads were assigned to *C. rosea* in the *C. rosea*-*F. graminearum* interaction. The summary data of transcriptome sequencing and mapping is presented in Table S5.

Compared with the *C. rosea* WT, the analysis identified 126 DEGs (106 upregulated and 20 downregulated) in Δ*dcl1* against *B*. *cinerea,* while this number was much higher against *F*. *graminearum* where 897 genes (504 upregulated and 393 downregulated) were differentially expressed (Table S6). Among these, a majority of genes were uniquely expressed in the respective interaction, as only 32 and three genes were commonly upregulated and downregulated, respectively, against both the mycohosts (Figure 3A). The deletion of *dcl2* affected the expression pattern of a higher number of genes compared to the deletion of *dcl1*. In Δ*dcl2*, in comparison to the WT, a total of 1894 (251 upregulated and 1643 downregulated) and 1706 (490 upregulated and 1216 downregulated) genes were differentially expressed against *B*. *cinerea* and *F*. *graminearum*, respectively (Table S6). In contrast to Δ*dcl1* where a relatively lower proportion of genes (15.7% against *B*. *cinerea*; 43.7% against *F*. *graminearum*) were downregulated, a higher proportion (87% against *B*. *cinerea*, 73% against *F*. *graminearum*) of DEGs in Δ*dcl2* were downregulated. Among the upregulated genes in Δ*dcl2,* 124 genes were commonly upregulated while 118 genes and 365 genes were uniquely upregulated against *B*. *cinerea* and *F*. *graminearum*, respectively. Among downregulated genes, 669 were common while 973 and 538 genes were unique against *B*. *cinerea* and *F*. *graminearum*, respectively (Figure 3B).

**Figure 3:**
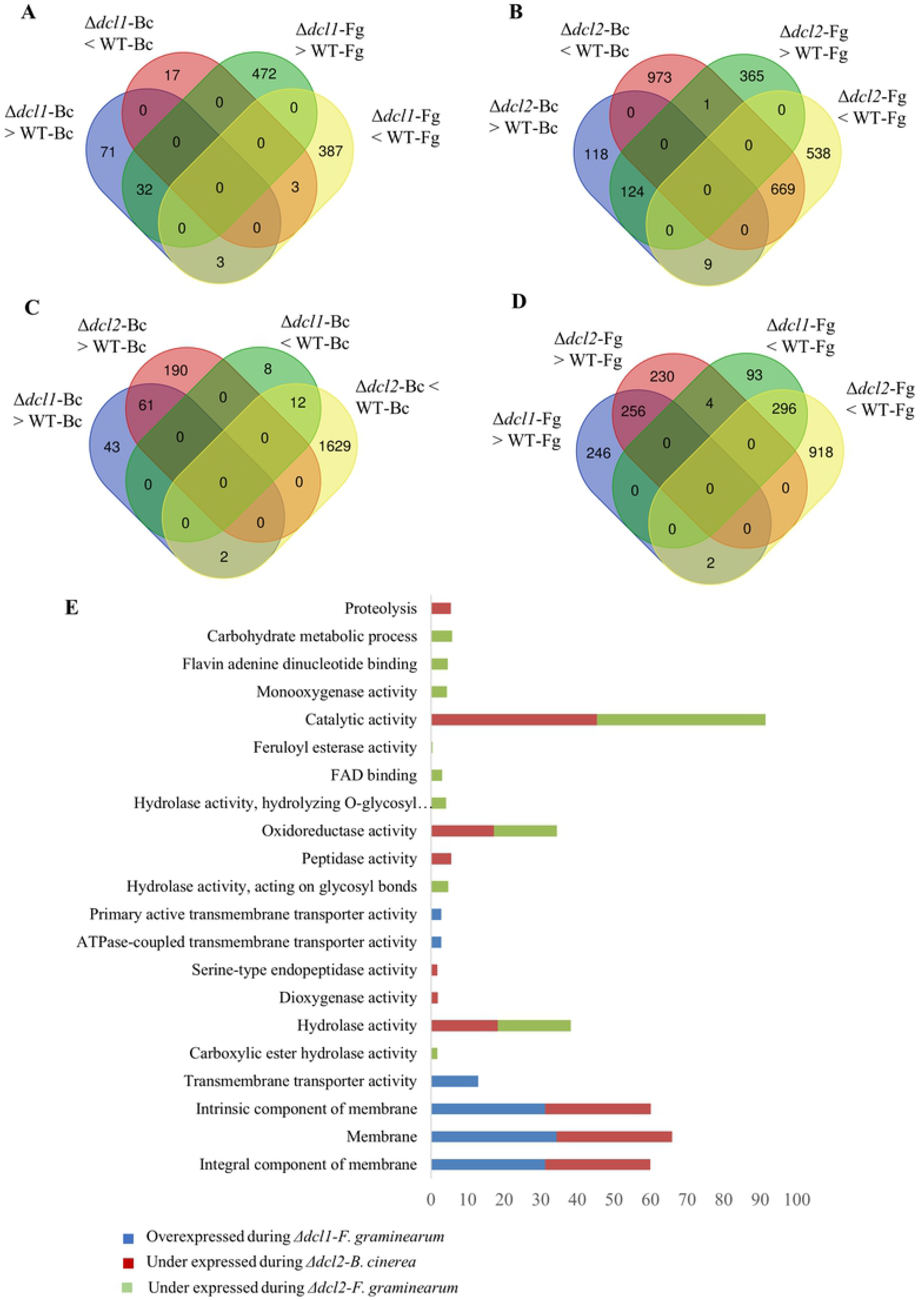
Transcriptome analysis of *C*. *rosea* WT and *dcl1* and *dcl2* deletion strains during the interactions with *B. cinerea* (Bc) and *F*. *graminearum* (Fg). **(A)** Venn diagram showing the common and species-specific DEGs in Δ*dcl1* against *B. cinerea* and *F. graminearum*; **(B)** Venn diagram showing the common and species-specific DEGs in Δ*dcl2* against *B. cinerea* and *F. graminearum*. (**C**) Overlap between DEGs genes in Δ*dcl1* and Δ*dcl2* against *B. cinerea* (**D**) Overlap between DEGs genes in Δ*dcl1* and Δ*dcl2* against *F. graminearum.* (**E**) Gene Ontology terms enriched in the differentially expressed *C. rosea* genes during the interactions.

The number of DEGs overlapping in Δ*dcl1* and Δ*dcl2* during the interactions with a common mycohost was determined (Figure 3C and 3D). Among the genes that were upregulated in Δ*dcl1* or Δ*dcl2 a*gainst *B*. *cinerea*, 61 were common, while 45 genes (41%) and 190 (76%) genes were uniquely upregulated in Δ*dcl1* and Δ*dcl2*, respectively. However, the number of genes downregulated in both mutants against *B. cinerea* was 12. During contact with *F. graminearum*, a similar number of genes was upregulated in the two mutants (246 in Δ*dcl1*, 230 in Δ*dcl2* and 256 in both), while the number of downregulated genes was greater in Δ*dcl2* (93 in Δ*dcl1*, 918 in Δ*dcl2* and 296 in both) (Figure 3C and 3D).

GO enrichment analysis was performed to evaluate which processes that were most affected in the *dcl* gene deletion mutants. Our results showed that a higher number of GO terms were significantly enriched in *C. rosea* genes under expressed in Δ*dcl2* compared to the whole transcriptome. In the molecular function category, we found that the terms such as catalytic activity (GO:0003824), hydrolase activity (GO:0016787) and oxidoreductase activity (GO:0016491) were commonly (against both the mycohosts) enriched (*P* ≤ 0.05) among the downregulated genes in Δ*dcl2*, indicating a role of these genes in mycoparasitism-related functions in *C*. *rosea* (Figure 3E). In contrast, other GO terms were only enriched against one of the two mycohosts. This was the case for the protein catabolism terms peptidase activity (GO:0008233) and proteolysis (GO:0006508), specifically enriched during the Δ*dcl2*-*B*. *cinerea* interaction. Carbohydrate metabolism-related terms like carbohydrate metabolism process (GO:0005975) and hydrolase activity acting on glycosyl bond (GO:0016798) were characteristic for the Δ*dcl2*-*F*. *graminearum* interaction (Figure 3E).

### 3.6 DCLs regulate genes with a predicted function during fungus-fungus interactions in *Clonostachys rosea*

Since the absence of DCL2 affected the production of secondary metabolites, antagonism and biocontrol of *C*. *rosea*, we performed an in-depth analysis of genes with a reported function during interspecific interactions in *C. rosea*, including membrane transporters, enzymes involved in biosynthesis of secondary metabolites and hydrolytic enzymes. Additionally, expression pattern of genes coding for transcription factors and various component of the silencing machinery were analyzed. For each of these categories, there were more upregulated genes than downregulated ones in the Δ*dcl1* strain. An opposite pattern was evident in the Δ*dcl2* strain, where the number of upregulated genes in each category tended to be higher than that of downregulated ones, except for ABC transporters (Table 1; supplementary table S7).

**Table 1:** Number of differential expression of genes in Δ*dcl1* and Δ*dcl2* compared with the WT during the *C. rosea* interaction with *F*. *graminearum* and *B*. *cinerea*.

|  | *C. rosea* - *F. graminearum* |  |  |  | *C. rosea* - *B. cinerea* |  |  |  |
| --- | --- | --- | --- | --- | --- | --- | --- | --- |
| | $\Delta dcl1$ | | $\Delta dcl2$ | | $\Delta dcl1$ | | $\Delta dcl2$ | |
|  | Up | Down | Up | Down | Up | Down | Up | Down |
| MFS transporters | 26 | 16 | 12 | 64 | 5 | 1 | 6 | 99 |
| ABC transporters | 14 | 0 | 10 | 6 | 1 | 0 | 4 | 3 |
| SM biosynthesis | 45 | 38 | 27 | 99 | 7 | 1 | 13 | 127 |
| Chitinases | 0 | 0 | 3 | 3 | 1 | 1 | 1 | 3 |
| Transcription factors | 24 | 6 | 31 | 28 | 5 | 1 | 17 | 56 |
| Gene silencing machinery | 4 | 0 | 4 | 1 | 0 | 0 | 1 | 3 |

#### 3.6.1 Membrane transporters

Deletion of *dcl2* affected expression of 161 major facilitator superfamily (MFS) transporters in *C*. *rosea*. Among these, 12 MFS transporters were upregulated and 64 were downregulated during the interaction with *F*. *graminearum*, while 6 were upregulated and 99 were downregulated during the interaction with *B*. cinerea (Table 1, supplementary table S7). Interestingly, 10 downregulated and 1 upregulated MFS transporters genes in Δ*dcl2* showed high sequence similarity (≥ 48 percent identity) with MFS transporters previously characterized for their involvement in efflux of secondary metabolites (polyketides, quinone and polyketide/non-ribosomal peptide hybrids) that are important for fungal virulence (Table 2). These included *apdF* (aspyridones efflux protein in *Colletothricum siamense*), *opS2* (quinone transporter in *Aspergillus udagawae*), *atB* (terreic acid efflux protein in *F. oxysporum*), *FUB11* (fusaric acid efflux pump in *Lachnellula suecica*), *FUBT* (efflux pump involved in export of fusaric acid in *F. culmorum*), *rdc3* (radicicol efflux pump in *F. oxysporum*) and *aflT* (aflatoxin efflux pump in *Phialocephala subalpine*) [102–105]. Furthermore, a homolog of *FUS6* (fusarin efflux pump FUS6 in *Colletothricum fructicola*) was upregulated. However, none of the corresponding gene clusters were present in the genome of *C. rosea*, suggesting that these MFS transporters constitute resistance proteins activated as a defense against harmful, hitherto unknown, secondary metabolites. Moreover, 22 MFS transporter genes were previously reported to be induced in *C*. *rosea* during the interactions with *B*. *cinerea* and *F*. *graminearum* [49]. Nine of these MFS transporter genes were significantly downregulated in the Δ*dcl2* strain during the interactions with *B*. *cinerea* or *F*. *graminearum* (Table 2). In summary the Δ*dcl2* mutant showed downregulation of transporters with predicted function in secondary metabolite export and putative detoxification.

**Table 2:** Differentially expression patterns of selected genes in *C. rosea* Δ*dcl1* and Δ*dcl2* strains during interactions with *B. cinerea* (Bc) or *F. graminearum* (Fg) compared to those of the *C. rosea* WT.

| Gene ID | Log2FC expression* |  |  |  | Comments |
| --- | --- | --- | --- | --- | --- |
| | $\Delta dcl1$-<br>Bc | $\Delta dcl1$-<br>Fg | $\Delta dcl2$-<br>Bc | $\Delta dcl2$-<br>Fg | |
| **Differentially expressed MFS transporters genes identical with previously characterized MFS transporters** |  |  |  |  |  |
| CRV2G00017900 | -0.36 | **-1.94** | 0.23 | **-5.05** | *mfs212* (ID 50% with apdF (PKS-NRPS transport)) |
| CRV2G00017824 | 0.36 | -0.68 | 0.21 | **-1.54** | *mfs* (ID 48% OpS2 (Quinone transport)) |
| CRV2G00015530 | -0.21 | -1.89 | 0.09 | **-2.28** | *mfs* (ID 59% with atB (terreic acid transport)) |
| CRV2G00015418 | 0.02 | **-1.61** | **-1.09** | **-1.56** | *mfs* (ID 60% with FUB11 (Polyketide transport)) |
| CRV2G00004817 | 0.53 | **-1.6** | **-4.04** | **-2.92** | *mfs506* (ID 57% with FUBT (Polyketide transport)) |
| CRV2G00002357 | -0.4 | -1.26 | **-1.69** | **-1.96** | *mfs533* (ID 70% with rdc3 (Polyketide transport)) |
| CRV2G00016200 | 0.12 | -0.69 | **-2.31** | **-2.18** | *mfs530* (ID 60% with rdc3 (Polyketide transport)) |
| CRV2G00004939 | 0.22 | -1.76 | **-2.09** | **-3.04** | *mfs534* (ID 80% with rdc3 (Polyketide transport)) |
| CRV2G00019617 | 1.94 | 4.06 | **1.59** | **3** | *mfs595* (ID 77% with FUS6 (Polyketide transport)) |
| CRV2G00011170 | 0.95 | 0.17 | 0.14 | **-3.32** | *mfs602* (ID 60% with aflT (Polyketide transport)) |
| CRV2G00005334 | 0.05 | -5.44 | **-4.55** | **-5.94** | *mfs589* (ID 70% with aflT (Polyketide transport)) |
| **Reduced expression of MFS transporters that were induced in *C. rosea* against *B. cinerea* or *F. graminearum*** |  |  |  |  |  |
| CRV2G00004685 | 0.32 | -0.79 | 0.62 | **-1.57** | *mfs464* |
| CRV2G00005389 | -0.81 | -0.75 | **-1.79** | **-1.38** | *mfs271* |
| CRV2G00018263 | -0.37 | -0.79 | -0.74 | **-2.14** | *mfs524* |
| CRV2G00011170 | -0.03 | -1.18 | 0.14 | **-3.32** | *mfs602* |
| CRV2G00012180 | 1.12 | **-2.65** | -1.45 | **-2.9** | *mfs166* |
| CRV2G00015972 | -0.06 | **-2.26** | **-1.77** | **-2.3** | *mfs205* |
| CRV2G00004853 | 0.45 | -1.45 | **-2.37** | **-2.27** | *mfs104* |
| CRV2G00004939 | 0.22 | **-1.76** | **-2.09** | **-3.04** | *mfs534* |
| CRV2G00018885 | -0.39 | -1.22 | **-3.55** | **-2.63** | *mfs24* |
| **Reduced expression of polyketide and non-ribosomal peptide synthetase genes** |  |  |  |  |  |
| CRV2G00011222 | -0.67 | 0.01 | 0.03 | **-1.88** | *pks14* |
| CRV2G00013582 | 0 | -1.43 | -0.03 | **-1.61** | *pks23* |
| CRV2G00015413 | 0.75 | **-2.28** | **-1.86** | **-2.96** | *pks12* |
| CRV2G00015415 | 1.09 | **-2.7** | **-3.22** | **-3.15** | *pks2* |
| CRV2G00018696 | -0.92 | -0.63 | -0.13 | **-4.97** | *pks6* |
| CRV2G00018222 | 0.03 | -1.43 | **-2.43** | **-1.79** | *pks22* |
| CRV2G00004952 | 0.11 | **1.88** | 0.74 | **1.54** | *nrps* |
| CRV2G00005605 | 0.65 | **2.73** | **1.95** | **2.33** | *nrps* |
| CRV2G00012656 | 0.18 | **1.82** | **1.95** | **2.17** | *nrps16* |
| CRV2G00015275 | -0.15 | -0.7 | 0.76 | **-2.06** | *nrps* |
| CRV2G00016915 | 0.67 | **-1.91** | **-3.07** | **-3.17** | *nrps* |
| CRV2G00014896 | 0.25 | 1.44 | 1.26 | **1.68** | *nrps9* |
| CRV2G00005211 | 0.26 | **-1.62** | **-3.74** | **-2.3** | Indole |
| CRV2G00002084 | **4.33** | 0.12 | **5.24** | -0.84 | Terpene |
| **Differentially expressed transcription factor genes identical with previously characterized transcription factors** |  |  |  |  |  |
| CRV2G00004759 | -0.69 | -0.32 | **-1.75** | -1.02 | ID 60% with FGR27 |
| CRV2G00006707 | -0.01 | -0.9 | **-1.62** | -1.31 | ID 73% with CCAAT-binding subunit HAP3 |
| CRV2G00015419 | 0.29 | -0.95 | **-2.22** | **-1.73** | ID 53% with sorbicillin regulator YPR2 |
| CRV2G00011734 | 0.32 | **1.81** | 0.56 | 1.41 | ID 79% with abaA |
| CRV2G00011385 | 0.19 | -0.46 | **2.58** | 1.16 | ID 57% with CTF1 |
| CRV2G00016352 | 0.73 | **1.51** | 0.47 | 1.3 | ID 65-70% SUC1 |
| CRV2G00019080 | **1.98** | **2.1** | 1.16 | **1.5** | ID 65% with SUC1 |
| CRV2G00019116 | 0.9 | **2.32** | 1.01 | **2.2** | ID 70% SUC1 |
| CRV2G00016935 | -0.74 | -0.22 | **-1.69** | -0.7 | ID 69% with prtT |
| CRV2G00018531 | -0.21 | -0.48 | **-2.12** | -1.35 | ID 61% with Sterol uptake control 2 |
| CRV2G00019093 | -0.38 | 0.43 | **-1.5** | -0.14 | ID 60% with GAL4 |
| **Differentially expressed Chitinases and N-acetylhexosaminidases genes** |  |  |  |  |  |
| CRV2G00001280 | -0.08 | -0.85 | **-3** | **-1.67** | Chitinase *ech42* |
| CRV2G00003425 | -0.3 | **-1.54** | **-3.6** | **-3.2** | Chitinase *ech37* |
| CRV2G00018858 | -0.01 | -0.06 | **-1.9** | **-1.82** | Chitinase *chia5* |
| CRV2G00017631 | -0.07 | 0.16 | 0.62 | **2.51** | Chitinase |
| CRV2G00006887 | 0.82 | **2.18** | 0.92 | **1.75** | Chitinase *ech58* |
| CRV2G00011101 | -0.3 | -0.13 | **2.25** | **2.1** | Chitinase *chic1* |
| CRV2G00002927 | -0.21 | -0.42 | **-1.76** | -0.78 | NAG |
| CRV2G00012950 | -0.14 | -0.43 | **-2.5** | **-2.26** | NAG |
| **Differentially expressed genes associated with gene silencing machinery** |  |  |  |  |  |
| CRV2G00000975 | 0.2 | 0.1 | 1.2 | **1.9** | Argonaute2-like |
| CRV2G00016556 | 0.2 | **2.1** | 0.4 | 1.3 | Chromatin remodelling protein |
| CRV2G00012165 | 0.2 | **4** | -0.4 | **4.3** | Histone deacetylase |
| CRV2G00007951 | 0.4 | 0.4 | 1 | **1.6** | Histone deacetylase |
| CRV2G00006603 | 0.9 | **2.3** | **2.4** | **2.3** | RNA helicase |
| CRV2G00007159 | 0.6 | **1.6** | 0.5 | 1 | RNA helicase |
| CRV2G00001612 | -0.6 | 0.1 | **-1.6** | **-1.8** | RNA helicase |
| CRV2G00012613 | -0.7 | 0.9 | **-2.4** | 0.1 | RNA helicase |
| CRV2G00009762 | 0 | 0.9 | **-1.7** | -0.6 | RNA-directed RNA polymerase |

\*Significant differences are highlighted in bold letters. FDR < 0.05 in combination with a log2FoldChange > 1.5 or < -1.5 was considered to define differentially expressed genes.

In contrast to the expression pattern of MFS transporters, a higher number of ATP-binding cassette (ABC) transporter genes was upregulated in both the deletion strains, but specifically against *F*. *graminearum* where 14 and 10 genes were upregulated in Δ*dcl1* or Δ*dcl2*, respectively (Table 1). Out of 19 ABC transporters that were differentially regulated in Δ*dcl2*, 5 upregulated and 1 downregulated belonged to the multidrug resistance protein (MDR) subfamily, 3 downregulated and 1 upregulated belonged to the multidrug resistance-associated protein (MRP) subfamily, and 4 upregulated and 1 downregulated belonged to pleiotropic drug resistance protein (PDR) subfamily (Table S7).

#### 3.6.2 Secondary metabolite biosynthetic genes

Genes associated with secondary metabolite production are often arranged in biosynthetic gene clusters (BGCs) that consist of genes coding for core enzymes typically non-ribosomal peptide synthetase (NRPS), polyketide synthase (PKS) or terpene cyclase, together with genes coding for additional proteins including modifying enzymes, transporters and transcription factors [106]. We used antiSMASH to predict the biosynthetic gene clusters in *C*. *rosea* and identified 33 NRPS BGCs, 29 PKS BGCs, 7 BGCs for terpenes and 7 for NRPS-PKS hybrids, and one BGC for indole and betalactone biosynthesis.

Gene expression analysis of both Δ*dcl1* and Δ*dcl2* identified a total of 230 DEGs predicted to be part of BGCs involved in secondary metabolite biosynthesis. Among the BGCs, the core biosynthetic genes in eight NRPS, five PKS, one terpene and one indole BGCs were differentially regulated in Δ*dcl2* against *B*. *cinerea* or *F*. *gramineaum* (Table 2; Table S7). Interestingly, NRPS and PKS BGCs core genes showed opposite expression pattern with each other as NRPS BGCs core genes were mostly upregulated in Δ*dcl2,* while PKS BGCs core genes were down regulated (Table 2). Among the downregulated core genes of PKS BGCs were the three PKS genes *pks22*, *pks2* and *pks12*, reported to be part of previously identified BGCs responsible for the production of clonorosein and sorbicillin, in *C*. *rosea* and *T*. *reesei*, respectively (Figure 4) [50, 107]. Sorbicillin is the precursor for sorbicillinol, which is in turn necessary for other sorbicillinoid compounds [108], explaining the low production of these substances by the Δ*dcl2* mutant.

**Figure 4:**
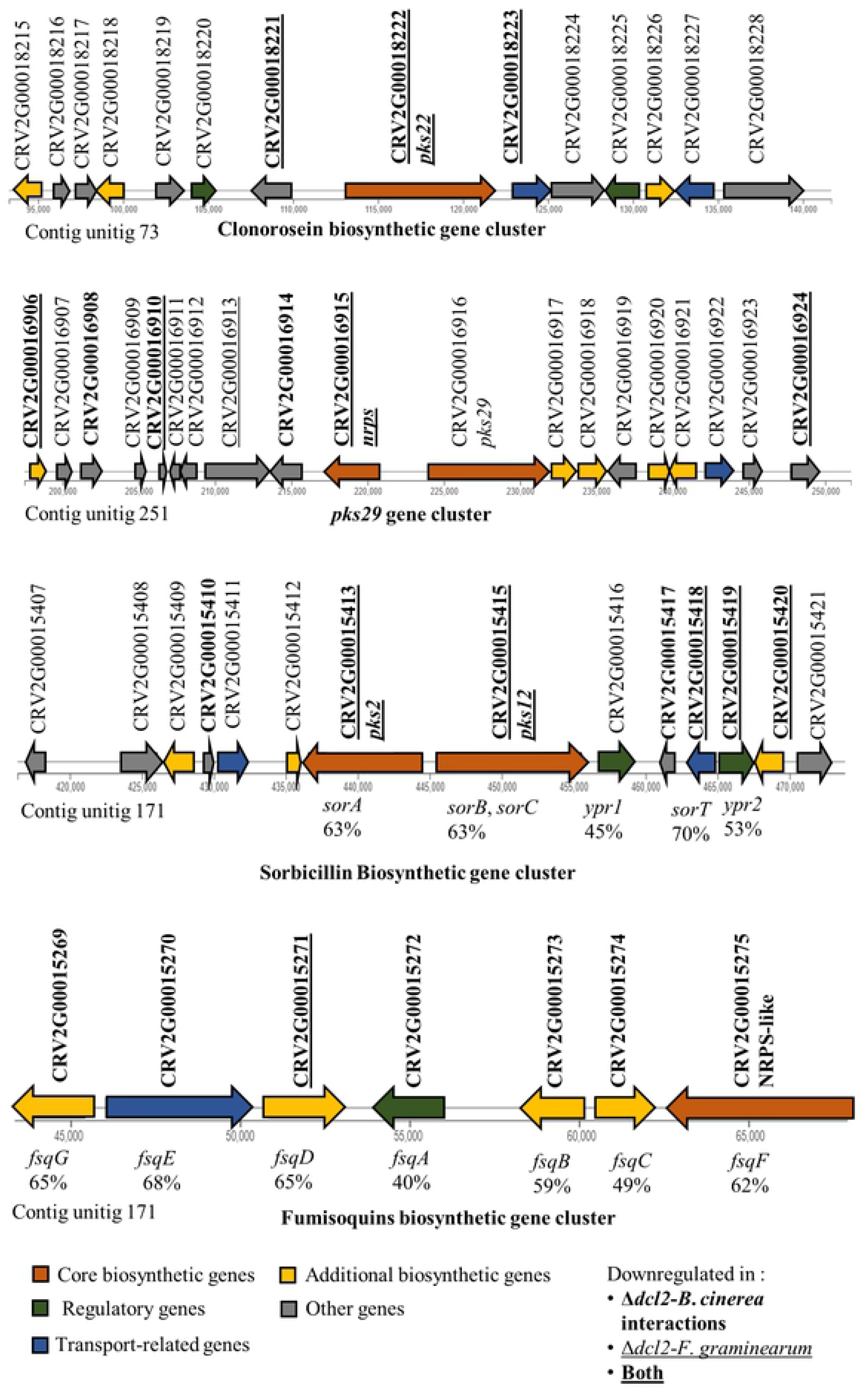
Expression of predicted *C. rosea* gene clusters of clonorosein, *pks29*, sorbicillin and fumisoquins. Gene IDs in bold letters indicate downregulated genes during *Δdcl2*-*B. cinerea* interactions; underline indicates downregulated genes during *Δdcl2*-*F. graminearum* interactions; Bold and underlined indicates genes that were downregulated against both the mycohosts. Bold and underlined indicates genes that were downregulated against both the mycohosts. The gene names for the sorbicillin and fumisoquin gene clusters were assigned through comparison with *Trichoderma reesei* and *Aspergillus fumigatus*, respectively [108, 118]. A minimum query coverage of 80% was required in the comparison and the maximum evalue was fixed at 1e-95.

#### 3.6.3 Transcription factors

The transcriptome analysis further identified 128 differentially expressed genes predicted to encode transcription factors in the Δ*dcl1* and Δ*dcl2* strains (Table 1, Table S7). We identified 11 transcription factors genes that were differentially expressed in Δ*dcl1* and/or in Δ*dcl2* and showed greater than 50% sequence identity with genes previously characterized for their role as transcriptional regulators. CRV2G00011734 was upregulated in the Δ*dcl1* and showed identity with the conidiophore development regulator gene *abaA* [109, 110], while CRV2G00016352, CRV2G00019080 and CRV2G00019116, also upregulated, showed identity with the sucrose metabolic gene *suc1*, shown to be associated with mitotic and meiotic cell division in fission yeast [111]. The genes CRV2G00004759, CRV2G00006707, CRV2G00015419, downregulated in Δ*dcl2*, showed identity with transcription factor genes *fgr27*, *hap3*, and *ypr2*, shown to be involved in regulating growth and secondary metabolite production [107,112,113] (Table 2). In summary, the Dicer-dependent control of transcription factor gene expression was to a large part mycohost-specific, with no transcription factors differentially expressed against both mycohosts in Δ*dcl2.* Moreover, among the identified transcription factors there were many homologs of genes known to have a role in regulating secondary metabolism and growth.

#### 3.6.4 Glycosyl hydrolase families 18 and 20

The *C*. *rosea* genome contains 13 genes coding for enzymes with predicted chitinase (glycoside hydrolase family 18 [GH18]) activity [44], six of which were differentially regulated in Δ*dcl2* against *B*. *cinerea* or *F*. *graminearum* (Table S7). Among these, CRV2G00001280 (*ech42*), CRV2G00003425 (*ech37*) and CRV2G00018858 (*chiA5*) were downregulated against both the mycohosts, while CRV2G00017631, CRV2G00006887 (*ech58*) and CRV2G00011101 *(chiC1*) were upregulated against both the mycohosts (Table 2). Furthermore, the *C*. *rosea* genome contains two genes (CRV2G00002927 and CRV2G00012950) coding for predicted N-acetylhexosaminidases (NAG; GH20), the expression of which was downregulated in Δ*dcl2* against *B. cinerea* (both genes) and *F. graminearum* (only CRV2G00012950). In summary, many glycoside hydrolases with a known role in degrading mycohost cell walls were downregulated in the Δ*dcl2* mutant after contact with the mycohosts.

#### 3.6.5 Genes associated with gene silencing machinery

To investigate an effect of *dcl1* and *dcl2* deletions on various protein components involved in the gene silencing machinery through chromatin modification in *C*. *rosea*, Blast2GO was used to identify genes encoding RNA helicases, chromatin remodeling proteins, histone deacetylases and histone methyltransferases. We identified 118 genes (excluding DCL, AGO and RDRP) including 67, 23, 18 and three genes coding for RNA helicases, chromatin remodeling proteins, histone deacetylases and histone methyltransferases, respectively (Table S8). Deletion of *dcl1* did not cause differential expression in the Δ*dcl1*-*B*. *cinerea* interaction, while during contact with *F. graminearum* we detected upregulation of two RNA helicase genes (CRV2G00006603, CRV2G00007159), one gene coding for a chromatin remodeling protein (CRV2G00016556) and a histone deacetylase gene (CRV2G00012172), while one histone deacetylase gene (CRV2G00012172) was downregulated (Table 2). During the Δ*dcl2*-B. *cinerea* interaction, one RNA helicase gene (CRV2G00006603) was upregulated, and two RNA helicases (CRV2G00001612, CRV2G00012613), as well as an RNA-directed RNA polymerase (CRV2G00009762) were downregulated. Conversely, during the Δ*dcl2-F*. *graminearum* interaction two histone deacetylases (CRV2G00012165, CRV2G00007951), one RNA helicase gene (CRV2G00006603) and one gene coding for an argonaute protein (CRV2G00000975) were upregulated, while one RNA helicase (CRV2G00001612) gene was downregulated. (Table 2). In summary, many genes involved in chromatin modification and gene silencing are affected by the deletion of the *dcl* enzymes, particularly *dcl2.* Most of these, including an argonaute protein, are upregulated, possibly due to the diminished presence of regulating sRNAs in the mutants.

### 3.7 Analysis of sRNAs characteristics in the *Clonostachys rosea* WT and the *dcl* deletion strains

To investigate the effect of sRNAs on transcriptional regulation in *C*. *rosea*, sRNA libraries from *C*. *rosea* WT, Δ*dcl1* and Δ*dcl2* strains interacting with *B*. *cinerea* or *F*. *graminearum* were sequenced. The sequencing produced 16 million reads per sample on average. Between 61% and 72% of these read pairs were composed of non-structural RNAs including rRNA, tRNA, snoRNA, snRNA, and were excluded from the further analysis. The remaining subset of reads that were 18-32 nt long were used for alignment to the genome of *C. rosea*, *B. cinerea* and *F. graminearum.* A summary of sRNA characteristics and their alignment to the respective genome is presented in table S9. sRNAs mapping exclusively to the *C*. *rosea*, *B. cinerea* or *F. graminearum* genome (unique sRNAs) were selected for further analysis. On average 42% of sRNA reads from *C*. *rosea*-*B. cinerea* interactions were aligned uniquely to one of the two organisms, while this percentage was only 18% for *C. rosea*-*F. graminearum* interactions. This is plausible because *C. rosea* is evolutionary closer to *F. graminearum* (both belong to order Hypocreales) than to *B. cinerea*.

We compared the characteristics of sRNAs produced in the Δ*dcl1* and Δ*dcl2* mutants to those of the WT. The analysis of length distribution showed a significant reduction in sRNAs with a size of 19-22 nt in the Δ*dcl2* compared with the WT, while no difference in sRNA abundance was found between the Δ*dcl1* and WT (Figure 5A). The analysis of the 5′ terminal nucleotide composition showed a reduced proportion of reads (27%) with 5’ end uracil (5’-U) in the Δ*dcl2* strain, compared with 32% −37% proportion of reads with 5’-U from the WT and Δ*dcl1* strains (Figure 5B). The origin of sRNAs was not significantly affected by the deletion of *dcl* genes, with most reads mapping to coding sequences (49%), followed by intergenic regions (25%), promoters (12.3%), 3’ UTRs (8%), introns (4%) and 5’ UTRs (1.5%). A higher proportion (83.5%) of sRNAs was mapped with sense orientation, rather than antisense one, similarly to what is reported in previous studies in *F*. *graminearum* and *T*. *atroviride* [20, 114], and this might be due to byproducts of mRNA degradation. However, the relative proportion of sRNAs mapping to the antisense direction was reduced from an average of 17.5% during WT-*B*. *cinerea* interaction to 14.3% during Δ*dcl2*-*B*. c*inerea* interaction (Table S9).

**Figure 5:**
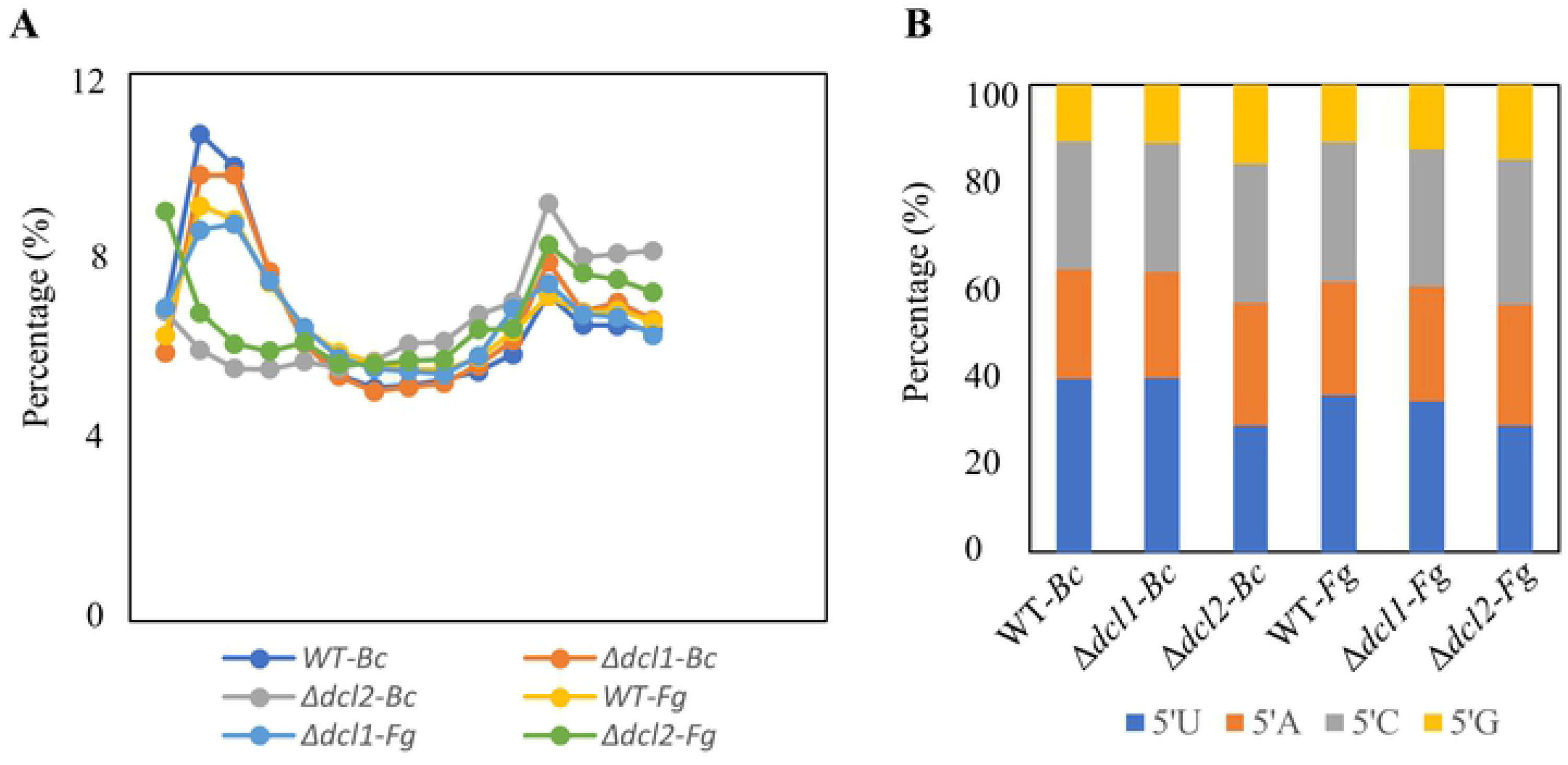
sRNAs characteristics in the *C. rosea* wild type (WT) and the *dcl* deletion strains. (**A**) Length distribution, and **(B**) 5’ end nucleotide preference of non-structural sRNAs produced by *C. rosea* WT and the *dcl* deletion strains during the interactions with *F. graminearum* (Fg) and *B. cinerea* (Bc). The only sRNAs comprised between 18 and 32 in length are considered.

#### 3.7.1 milRNA prediction in *Clonostachys rosea*

Mirdeep2 analysis predicted 61 milRNAs in *C*. *rosea* with a length between 18 and 25 nt and they were named cro-mir. These milRNAs originated from a variety of positions in the genome including promoters, introns, CDSs and UTRs, but mainly (28 of 61) from intergenic regions (Table S10). The expression of 15 cro-mirs was common against both the mycohosts, while 29 and 17 cro-mirs were expressed specifically during the interaction with *B. cinerea* or *F. graminearum*, respectively (Table S10). Interestingly, no cro-mir was found to be differentially expressed in Δ*dcl1* during the interspecific interactions, while 11 cro-mirs were significantly downregulated in Δ*dcl2* during the interaction with both mycohosts (Table 3). This downregulation was confirmed through stem-loop RT-qPCR (Table 3). A single milRNA (cro-mir-23) was identified as upregulated in the Δ*dcl2* mutant in the RNAseq analysis, but downregulated according to stem-loop RT-qPCR.

**Table 3:**
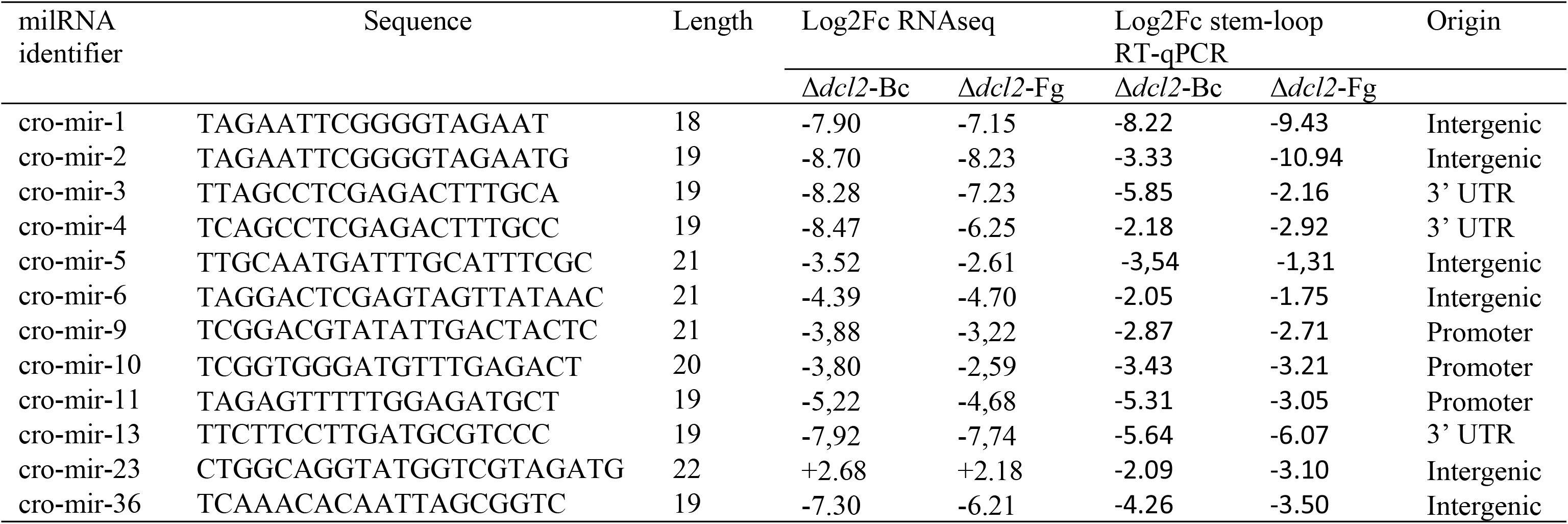
Differentially expressed *C*. *rosea* milRNAs, their length and locus of origin.

#### 3.7.2 Identification of cro-milRNAs endogenous gene targets

Twenty-one putative endogenous gene targets were identified for the 11 cro-mirs downregulated in Δ*dcl2* (Table 4). Eight gene targets were commonly upregulated in Δ*dcl2* during the interaction with *B*. *cinerea* and *F. graminearum*, while seven and six gene targets were uniquely upregulated during the interactions with *B. cinerea* and *F. graminearum*, respectively (Table 4). Among the predicted gene targets, several had putative regulatory roles:

**Table 4:** Endogenous putative gene targets in *C. rosea*, their expression pattern and predicted function.

| miRNA identifier | Gene target | Expression Log2Fc* |  | Target gene family | Characterized/putative function |
| --- | --- | --- | --- | --- | --- |
| | | $\Delta dcl2$-Bc | $\Delta dcl2$-Fg | | |
| cro-mir-3 | CRV2G00002264 | **1.08** | **1.42** | Serine/threonine-protein kinase (Gin4) | Septin ring assembly, Intracellular signal transduction |
| cro-mir-5 | CRV2G00013335 | **1.39** | **1.25** | Unknown | Unknown function |
| cro-mir-5 | CRV2G00015277 | **2.54** | **3.52** | Transcription factor | 60 S ribosome biogenesis |
| cro-mir-10 | CRV2G00015277 | **2.54** | **3.52** | Transcription factor | 60 S ribosome biogenesis |
| cro-mir-11 | CRV2G00015277 | **2.54** | **3.52** | Transcription factor | 60 S ribosome biogenesis |
| cro-mir-13 | CRV2G00001868 | **1.95** | **2.72** | Helicase | Chromatin re-modelling |
|  | CRV2G00002266 | **1.81** | **1.98** | Transcriptional regulator prz1 | Regulates the expression of Pmc1 ATPase Ca2+ pump |
| cro-mir-36 | CRV2G00013380 | **2.42** | **3.36** | ATPase | ATPase activity |
|  | CRV2G00005499 | **1.38** | **1.8** | Unknown | Unknown function |
|  | CRV2G00000111 | **1.95** | **2.69** | Unknown | Unknown function |
|  | CRV2G00014914 | **1.21** | 0.82 | Oxidation-reduction process | Part of secondary metabolite BGC |
| cro-mir-1 | CRV2G00003756 | **1.06** | 0.89 | tRNA ligase | Protein biosynthesis |
| cro-mir-2 | CRV2G00003756 | **1.06** | 0.89 | tRNA ligase | Protein biosynthesis |
| cro-mir-3 | CRV2G00008014 | **1.12** | 0.23 | GTPase-activating protein 2 | Signal transduction |
| cro-mir-6 | CRV2G00002043 | **1.12** | 0.99 | Transcription factor | Regulation |
| cro-mir-3 | CRV2G00009307 | **1.26** | 0.81 | Sterol O-acyltransferase 2 | Cholesterol metabolic process |
| cro-mir-11 | CRV2G00009307 | **1.26** | 0.81 | Sterol O-acyltransferase 2 | Cholesterol metabolic process |
| cro-mir-3 | CRV2G00011242 | **1.26** | 0.75 | Oxidoreductase | Oxidation-reduction |
| cro-mir-4 | CRV2G00011242 | **1.26** | 0.75 | Oxidoreductase | Oxidation-reduction |
| cro-mir-13 | CRV2G00004332 | **1.06** | 0.43 | GTP-binding protein | Ribosome biogenesis |
| cro-mir-1 | CRV2G00005300 | 0.69 | **1.38** | Unknown | Unknown function |
| cro-mir-4 | CRV2G00004339 | 0.48 | **1.03** | SNF2 RNA helicase | Chromatin re-modelling |
| cro-mir-9 | CRV2G00004339 | 0.48 | **1.03** | SNF2 RNA helicase | Chromatin re-modelling |
| cro-mir-10 | CRV2G00004339 | 0.48 | **1.03** | SNF2 RNA helicase | Chromatin re-modelling |
| cro-mir-11 | CRV2G00000903 | 0.82 | **1.03** | Unknown | Unknown function |
| cro-mir-36 | CRV2G00000903 | 0.82 | **1.03** | Unknown | Unknown function |
| cro-mir-10 | CRV2G00011823 | 0.93 | **1.21** | Choline-sulfatase | hydrolase activity |
| cro-mir-36 | CRV2G00011823 | 0.93 | **1.21** | Choline-sulfatase | hydrolase activity |
| cro-mir-4 | CRV2G00012062 | -0.18 | **1.09** | Unknown | Unknown function |
| cro-mir-13 | CRV2G00012781 | 0.3 | **1.01** | Unknown | Unknown function |

\* Upregulated (FDR < 0.05 in combination with Log2Fc >1) gene targets are highlighted in bold.

CRV2G00015277, CRV2G00002266 and CRV2G00002043 were predicted to encode putative transcription factors, CRV2G00001868 encodes an ATP-dependent helicase, while CRV2G00004332 and CRV2G00008014 encode a GTP binding protein and a GTPase with a putative role in signal transduction. Moreover, CRV2G00014914 was located in a secondary metabolite gene cluster and might have a role in regulating secondary metabolism (Table 4).

#### 3.7.3 Cross-species gene target identification

Using the criteria described for the endogenous gene target prediction, we identified 513 putative cross-species gene targets in *B*. *cinerea* (Table S11). Among these, the seven genes *bcpls1*, *bcpka1*, *bcnoxA*, *bcste11*, *bccap9*, *bccrh1* and *bcchsIV* were previously characterized for their role in growth and development, proteolysis and consequently in virulence (Table 5). Moreover, a gene encoding a *B. cinerea* homolog of SSAMS2 (BCIN_08g03180) was also among the putative targets, and this gene encodes a GATA transcription factor required for appressoria formation and chromosome segregation in *Sclerotinia sclerotiorum* [115]. Additionally, *bcnog1* and *bchts1* encoding for proteins putatively involved in ribosome biogenesis, and *bcphy2* and *bchhk1* encoding for signal transduction proteins were also identified as putative targets. Finally, three genes coding for a protein with a putative role in chitin recognition (*bcgo1*), chromatin remodeling (*bcyta7*) and intracellular trafficking and secretion (*bcvac8*) were also identified (Table 5).

**Table 5:**
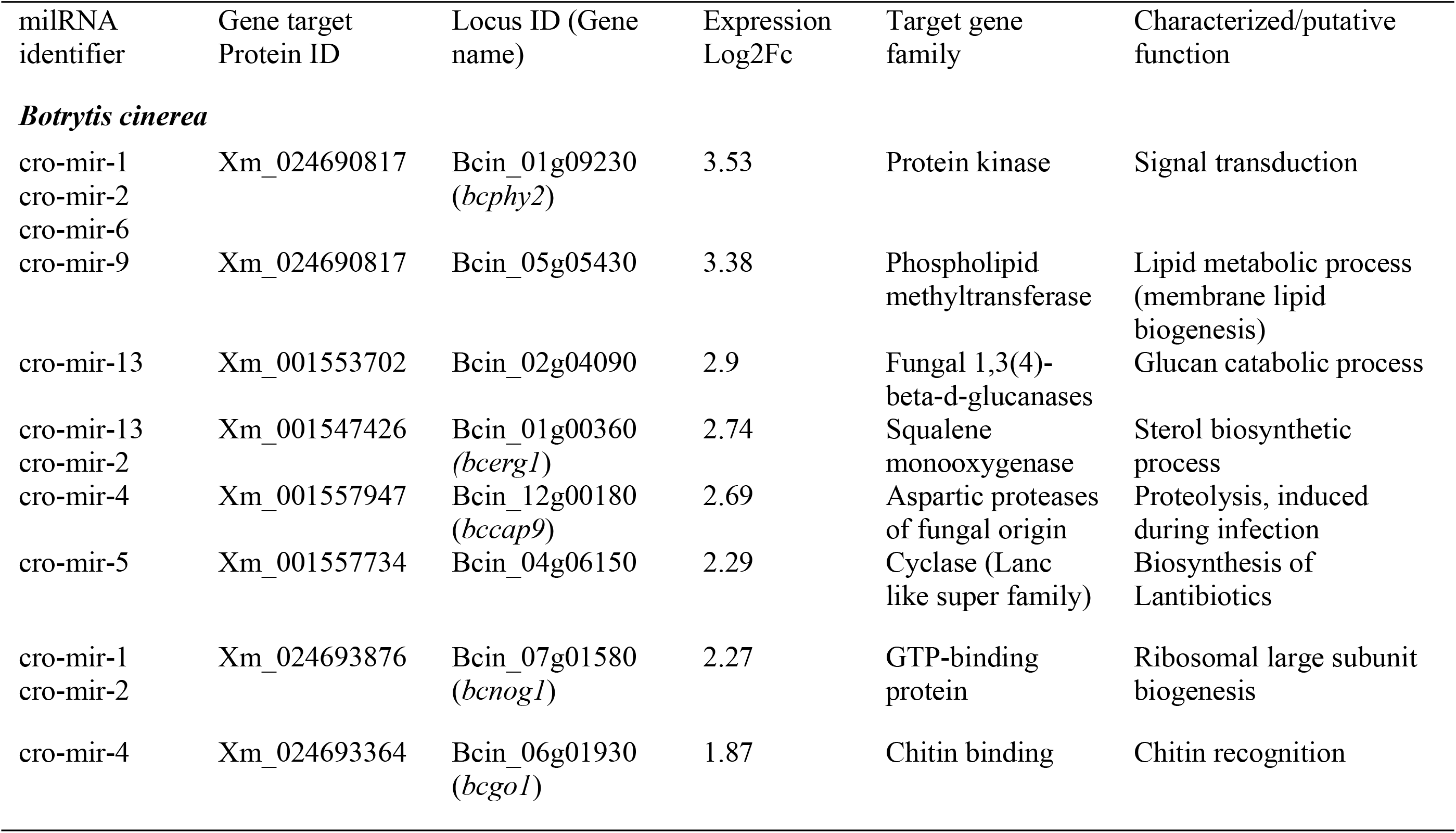
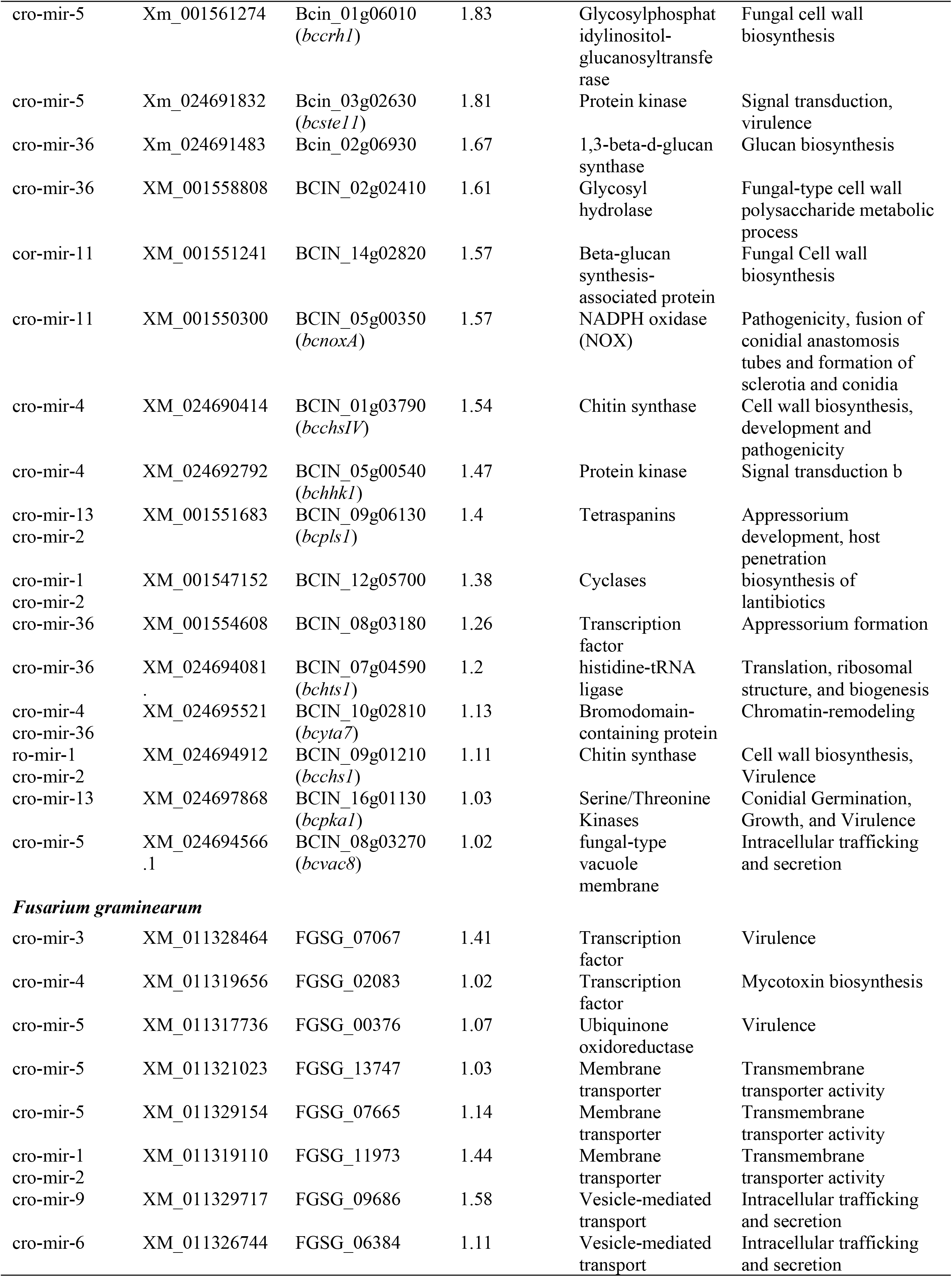
Most important cross-species putative gene targets in *B*. *cinerea* and *F*. *graminearum*, their expression pattern and putative function.

Thirty-five cross-species gene targets were predicted in *F*. *graminearum* as well. We identified three previously characterized virulence factors (FGSG_07067, FGSG_02083 and FGSG_00376) as putative targets of cro-mir-3, cro-mir-4, and cro-mir-5, respectively (Table 5). In addition, three membrane transporter genes (FGSG_13747, FGSG_13747, FGSG_13747) and two genes coding for proteins with a putative role in intracellular trafficking and secretion (FGSG_09686 and FGSG_09686) were identified as putative targets (Table 5). In summary, several mycohosts genes with a role in virulence, intracellular trafficking, secretion and regulation were identified as putative targets of *C. rosea dcl2*-dependant milRNAs.

### 3.8 Botrytis cinerea and Fusarium graminearum responded differently towards *Clonostachys rosea* WT and *dcl* deletion strains

Transcriptome analysis of *B*. *cinerea* and *F*. *graminearum* was performed to investigate whether deletion of *dcl* genes affects their response mechanism towards *C*. *rosea*. Read pairs unique to *B*. *cinerea* from the *C*. *rosea*-*B. cinerea* interaction, and unique to *F*. *graminearum* from the *C. rosea*-*F*. *graminearum* interaction were used in the analysis. From the total amount of read pairs that originated from the *C. rosea*-*B. cinerea* or *C*. *rosea*-*F*. *graminearum* interactions, 25% and 23% reads were uniquely assigned to *B. cinerea* and *F. graminearum*, respectively (Table S5).

In comparison to the WT-*B. cinerea* interaction, 24 genes (21 upregulated and three down-regulated) were differentially expressed in *B. cinerea* during the Δ*dcl1-B. cinerea* interaction. However, 721 genes were found to be differentially regulated (655 upregulated and 66 down) in the interaction with Δ*dcl2* (Figure 6A; Table S11). The 21 *B*. *cinerea* genes that were upregulated against Δ*dcl1* were also upregulated against Δ*dcl2* (Figure 6 A). We specifically investigated genes coding for hydrolytic enzymes, transcription factors, membrane transporters, known virulence factors, RNA silencing component proteins, and genes part of secondary metabolite BGCs. During the Δ*dcl1*-*B*. *cinerea* interaction, one gene (BCIN_14g03930) coding for a known virulence factor and two genes coding for MFS transporters were upregulated, while two genes that were part of secondary metabolite BGCs were down-regulated in *B*. *cinerea*. Deletion of *dcl2* induces increased expression of 12 genes previously characterized for their role in growth and development, virulence and pathogenesis in *B. cinerea*. Among the other genes, we detected the upregulation of GTPases, kinases, chitinases, squalene monooxygenases, and genes involved in chitin synthesis and chitin recognition (Table 6).

**Figure 6:**
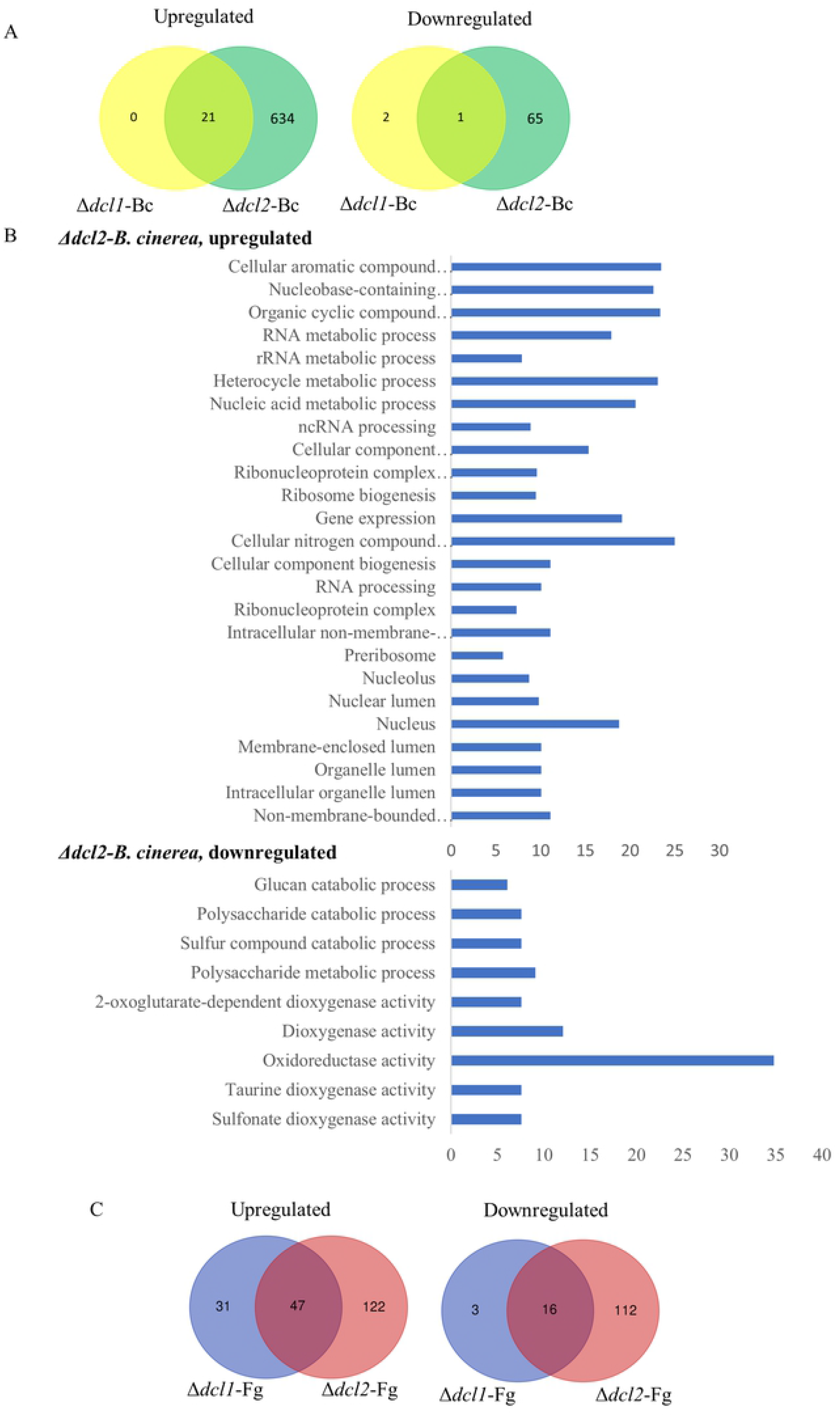
Transcriptome analysis of *B*. *cinerea* (Bc) and *F*. *graminearum* (Fg) during the interaction with *dcl1* and *dcl2* deletion strains compared to those of the WT. **(A)** Venn diagrams showing the overlap between upregulated and down regulated genes in the Δ*dcl1* and Δ*dcl2* strains during the interactions with *B. cinerea* compared with the WT. **(B)** Gene Ontology terms of enriched upregulated and downregulated gene in *dcl2* deletion strains during the interactions with *B. cinerea*. **(C)** Venn diagrams showing the overlap between upregulated and down regulated genes in the Δ*dcl1* and Δ*dcl2* strains during the interactions with *F. graminearum* compared the WT.

**Table 6:**
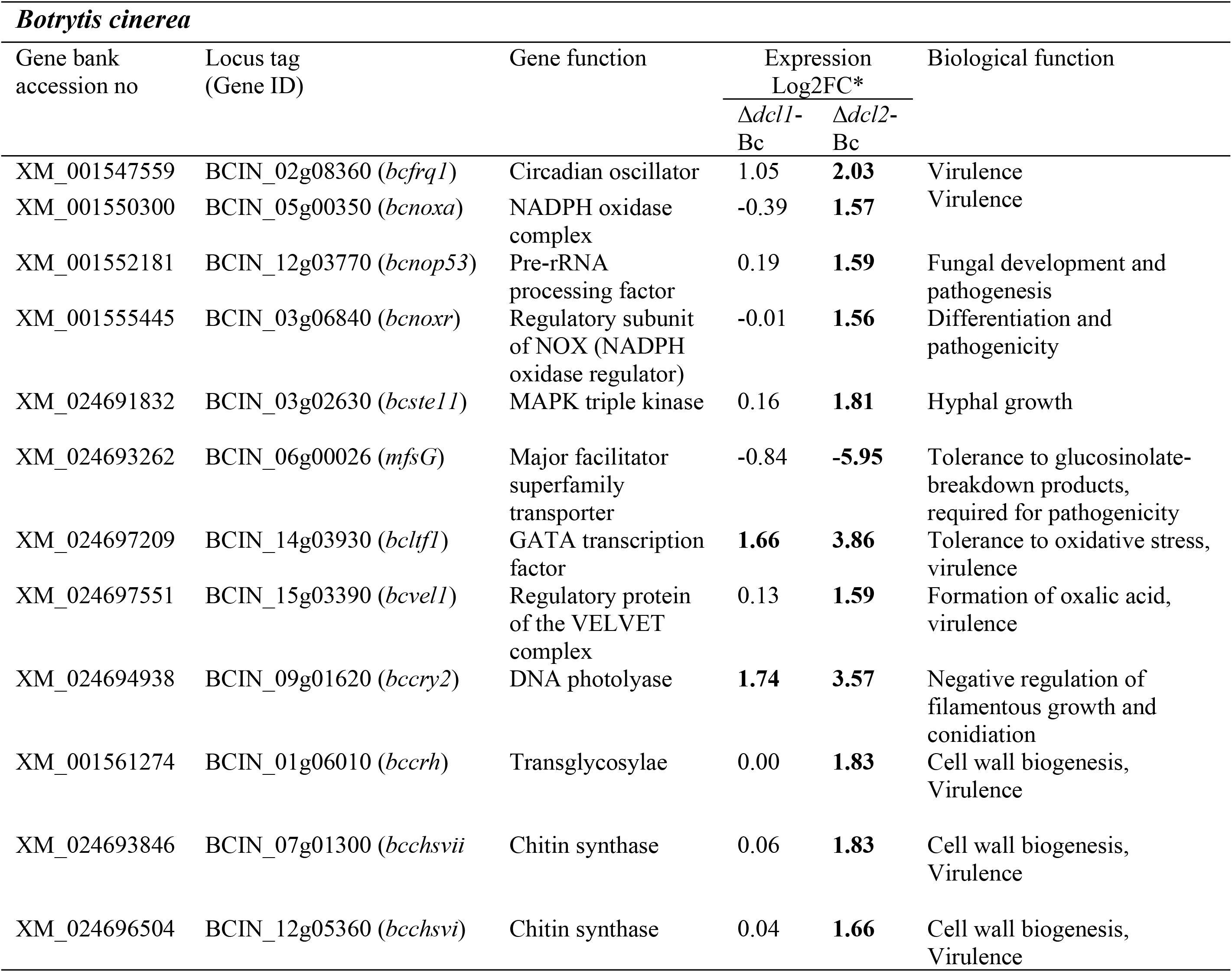
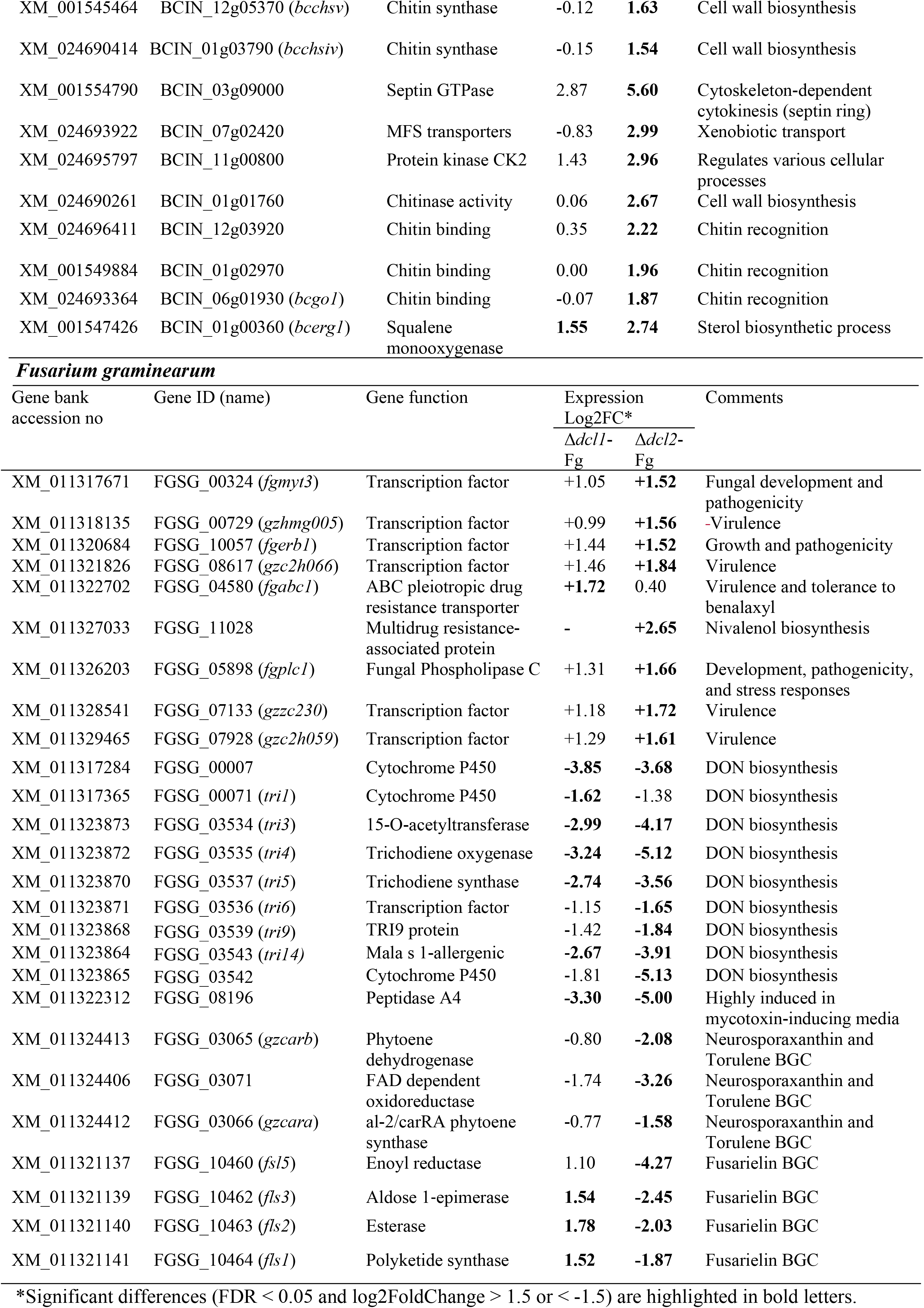
Differentially expression patterns of selected genes in *B*. cinerea and *F*. *graminearum* during the interaction with Δ*dcl1* and Δ*dcl2* compared to those of the *C. rosea* wild type and the same mycohost.

The other differentially expressed genes did not have a characterized functional role, but a function was predicted for some of them. In particular, among the genes upregulated during the Δ*dcl2*-*B*. *cinerea* interaction, we detected 49 putatively coding for hydrolytic enzymes, 24 located in putative secondary metabolite BGCs, 22 transcription factors, 17 genes involved in RNA silencing, 15 protein kinases and 13 MFS transporters (Table S11). GO enrichment analysis of upregulated genes during the Δ*dcl2*-*B*. *cinerea* interactions identified terms for metabolic processes, including gene expression (GO:0010467), cellular component organization or biogenesis (GO:0071840) and RNA processing (GO:0006396) (Figure 6B). However, GO terms oxidoreductase activity (GO:0016491), oxidation-reduction processes (GO:0055114), polysaccharide and glucan catabolic processes (GO:0000272 and GO:0009251) were enriched for the downregulated genes (Figure 6B).

During the Δ*dcl2-F. graminearum* interaction, 397 (169 upregulated and 128 downregulated) *F*. *graminearum* genes were differentially expressed, while only 97 (78 upregulated and 19 downregulated) were differentially expressed during the Δ*dcl1-F*. *graminearum* interaction (Figure 6C; Table S12). Forty-seven and 16 genes were respectively upregulated and downregulated against both mutant strains, while the rest were differentially expressed only during contact with one of the mutants. (Figure 6C). Furthermore, we found 26 (nine upregulated and 17 downregulated) previously characterized *F*. *graminearum* genes that were differentially regulated during the interaction with *dcl* deletion strains compared to the WT (Table 6). The downregulated genes included several ones involved in deoxynivalenone, neurosporaxanthin, torulene and fusarielin biosynthesis. Moreover, eight of the nine upregulated genes were previously characterized for having a role in *F. graminearum* virulence, and six of them encoded transcription factors (FgMYT3, GzHMG005, FgERB1, GzC2H066, GzZC230, GzC2H059) (Table 6). Moreover, during the interaction with the Δ*dcl2* mutant, 14 *F. graminearum* CAZyme genes showed upregulation with respect to the WT, all of them predicted to encode glycoside hydrolases, while only three were downregulated. MFS transporters were among the DEGs as well, with five of them being upregulated while seven were downregulated (Table S12).

## 4. Discussion

While the Δ*dcl1* mutant had a phenotype largely similar to the WT, the Δ*dcl2* mutant displayed evident differences, including a higher number of differentially expressed genes during the interaction with the plant pathogenic mycohosts. This amount of DEGs was significantly higher than the number of genes predicted to be directly targeted from DCL2-regulated milRNAs, but it has already been observed in *F. graminearum* and *T. atroviride* how RNA interference can be involved in regulating the activity of transcription factors and other regulatory elements, therefore indirectly influencing the expression of a vast array of genes and pathways [20, 114]. In our dataset, we could observe four *C. rosea* transcription factors downregulated in the WT during interaction with the mycohosts and putatively targeted by milRNAs downregulated in the Δ*dcl2* mutants. Among these, CRV2G00015277 and CRV2G00002266 were involved in the interaction with both the mycohosts, while CRV2G00002043 and CRV2G00000111 were involved only in response to *B. cinerea*. CRV2G00002266 exhibited significant sequence similarities with the PRZ1 transcription factor, known for regulating the expression of the vacuolar ATPase Ca2+ pump PMC1 [116]. This pump shown to regulates the level of cytoplasmic Ca2+ by activating Ca2+ dependent enzymes involved in protein secretion in the nuclear envelope, endoplasmic reticulum, golgi complex and trans-golgi/endosomal network in *S. cerevisiae* [116].

Furthermore, several other putative milRNA targets could have regulatory roles, including the predicted helicases CRV2G00001868 and CRV2G00004339 and putative Rho-type GTPase activating protein CRV2G00008014. In particular, the transcript of gene CRV2G00004339, putatively targeted by milRNAs during interaction with *F. graminearum*, encodes a helicase of superfamily SNF2, involved in cromatin remodeling by deposition of H2A [117].

Beyond the direct action of milRNAs on targets, the deletion of *dcl1* and especially *dcl2* induced the differential expression of several secondary metabolite BGCs in *C. rosea*. The BGC containing the PKS gene *pks22*, involved in the synthesis of the antifungal compound clonorosein [50] was downregulated in Δ*dcl2* during the interaction with both mycohosts. In contrast, no difference in clonorosein A production was detected between the WT and the *dcl* mutants in the metabolome analysis. However, as the metabolome analysis was performed during *in vitro* conditions, it is possible that the *dcl2*-dependent regulation of clonorosein production is more pronounced during contact with the mycohosts. In fact, *pks22* was previously shown to be induced during interactions with *B*. *cinerea* and *F*. *graminearum* [50]. The sorbicillin BGC, responsible for the yellow coloration of WT *C. rosea* colonies [50], is downregulated in Δ*dcl2*, and both sorbicillin and sorbicillinol were underproduced in Δ*dcl2* and had their biosynthesis restored in the complementation mutant in the *in vitro* trials, explaining the difference in pigmentation of the Δ*dcl2* mutant. This gene cluster was also induced during the interaction of *C. rosea* strain ACM941 with *F. graminearum* in the study of Demissie et al. [48]. However, it is interesting to notice how the positive regulator of the cluster, YPR1 (CRV2G00015416), is not differentially expressed in our study, while the transcription factor YPR2 (CRV2G00015419) is down-regulated and hence coregulated with the other genes in the gene cluster in Δ*dcl2*. YPR2 is a Gal4-like transcription factor predicted to positively regulate a negative regulator of sorbicillin biosynthesis [107], and its coregulation with the biosynthetic genes suggests that the deletion of DCL2 affects the control of sorbicillin production at a currently unknown level.

Furthermore, two putatively important BGCs were specifically downregulated in the Δ*dcl2* mutant during contact with *F. graminearum*: these were the *pks29* BGC involved in antagonism and biocontrol [50], and the BGC with the NRPS-like CRV2G00015275 as core enzyme. This last cluster was studied as “Cluster 3” in the work of Demissie et al. [47], where it was found to be induced in *C. rosea* after exposure to the *F. graminearum* secretome, and it presents strong homology with the fumisoquin cluster of *Aspergillus fumigatus* [118]. Deletion of the core NRPS-like enzyme of the cluster leads to reduced growth and sporulation in *A. fumigatus* [119], but fumisoquins were not produced in detectable amounts either by the WT or by the Δ*dcl2* mutant in our *in vitro* analysis. Biosynthesis of the corresponding compound in *C. rosea* may be specifically triggered during contact with *F. graminearum*. The transcription factor CRV2G00015277, putatively targeted by DCL2-dependant novel milRNAs cro-mir-5, cro-mir-10 and cro-mir-11, is located next to the cluster, and is upregulated in the Δ*dcl2* mutant. It is possible that CRV2G00015277 is a negative regulator of the cluster, targeted by milRNAs to induce the production of fumisoquins, but this hypothesis should be examined in a future study. None of these gene clusters (sorbicillin, clonoroseins, *pks29*, fumisoquins) were downregulated in the Δ*dcl1* mutant. The reduced production of bisorbicillinol in the Δ*dcl2* mutant also suggests that the deletion might hamper this fungus’ antibacterial properties, as several bisorbicillinoids synthesized by *C. rosea* have significant antibacterial activity [120].

A further reason for the diminished capacity of the Δ*dcl2* mutant to control the plant pathogenic mycohosts can be found in the downregulation of genes encoding enzymes involved in the degradation of the fungal cell wall. In the Δ*dcl2* mutant, between 55 and 64 glycoside hydrolase genes were downregulated compared to the WT. Among these were three GH18 chitinases (*ech37, ech42* and *chiA5*) and one GH20 N-acetylhexosaminidase (CRV2G00012950), which were downregulated during interaction with both mycohosts. Furthermore, four genes putatively involved in cell wall degradation of *F. graminearum* [48] were found to be downregulated in the Δ*dcl2* mutant: these were two glycoside hydrolases of classes GH2 (CRV2G00016896) and GH114 (CRV2G00003509), as well as two metallopeptidases (CRV2G00010851 and CRV2G00011092). Interestingly, the gene *chiC1,* predicted to encode a killer toxin-like chitinase that permeabilizes the cell wall of antagonistic species to facilitate entry of toxic metabolites [121, 122], is upregulated in the Δ*dcl2* mutant. This may be explained by the fact that *chiC1* is induced by chitin [44] and that the Δ*dcl2* mutant is compromised in its ability to antagonize the mycohosts, resulting in higher amounts of chitin exposed to the Δ*dcl2* mutant.

Moreover, 17 genes upregulated during *C. rosea* response to mycohosts in the study of Nygren et al. [49] were downregulated in the Δ*dcl2* mutants in comparison with the WT upon contact with the same mycohots. Among them is a putative isotrichodermin C-15 hydroxylase (*cyp1*), a type of protein also induced during mycoparasitism in *T. cf. harzianum* [123], but the majority of these genes is constituted by transporters, especially MFS transporters. This group includes gene *mfs464*, suggested in the study of Nygren et al. [49] to perform an important function in the mycoparasitic attack against *F. graminearum*, due to its extreme induction (fold change > 693). *Mfs166* and *mfs464*, downregulated in Δ*dcl2*, were found to be upregulated during *C. rosea* response to *F. graminearum* in the studies of both Nygren et al. [49] and Demissie et al. [48], making their involvement in response to the mycohot very likely. The other detected differentially expressed MFSs are commonly involved in efflux-mediated protection against exogenous or endogenous secondary metabolites and sugar uptake, suggesting a DCL-dependent influence on this aspect of *C. rosea* mycoparasitic action. This group also includes nine genes belonging to the Drug: H + Antiporter-2 family, which underwent a significant gene expansion during *C. rosea* evolution and has therefore a putative important role in the fungus lifestyle [124]. DCL-based control of these transporters is most likely indirect, because most MFS genes detected in this way are downregulated in the mutants, while direct targets of RNA silencing are expected to be upregulated after *dcl* deletion. Reinforcing this hypothesis, none of the MFS transporters predicted in *C. rosea* is a putative target of differentially expressed milRNAs detected in this study. Identification of several upregulated genes coding for MFS transporters used by mycohosts to tolerate harmful secondary metabolites of their own production strengthens the hypothesis that these proteins enable *C. rosea* to withstand mycohost-produced toxins during fungal-fungal interaction.

The differential expression of this vast amount of genes is likely due to the 128 putative transcription factors differentially expressed in the Δ*dcl2* mutant. Among these, CRV2G00006707 is homologous of the CCAAT-binding subunit HAP3, regulating growth and secondary metabolism in other filamentous fungi such as *F. verticillioides* [113, 125]. This gene is downregulated in the Δ*dcl2* mutant during interaction with both mycohosts (log2FC of −1.6 in Cr-Bc and −1.3 in Cr-Fg). Another transcription factor downregulated in the Δ*dcl2* mutant was CRV2G00004759, a homolog of the filamentous growth regulator 27 (*fgr27*) of *Trichoderma lentiforme*, which is involved in adherence regulation and could have a role in reduced growth rate of the mutant [112]. Moreover, two putative homologs of the sucrose utilization protein 1 (SUC1) are upregulated in the Δ*dcl2* mutant and its upregulation is associated to a delay in mitotic and meiotic nuclear divisions in *Schizosaccharomyces pombe* [111].

It is possible that part of the reduced ability of the Δ*dcl2* to overgrow *B. cinerea in vitro* and control *F. graminearum in vivo* comes from a cross-regulating action of *C. rosea* milRNAs targeting mycohost genes involved in the development or reduction of virulence. Specifically, three *F. graminearum* virulence factors were both downregulated during interaction with the WT *C. rosea* and putatively targeted by milRNAs downregulated in the Δ*dcl2* mutants. These genes included FGSG_07067, the GzZC232 transcription factor whose deletion impaired virulence in the work of Son *et al.* [126], FGSG_00376, the NOS1 NADH ubiquinone oxidoreductase proven to be a factor of virulence by Seong *et al.* [127] and FGSG_02083, the transcription factor ART1, whose deletion causes reduced starch hydrolysis and virulence, as well as the incapability of trichothecenes biosynthesis [128]. Regarding *B. cinerea*, among the putative milRNA-targeted downregulated genes, there were those encoding BCIN_09g06130, the BcPls1 tetraspanin necessary for appressorium-mediated penetration into host plant leaves [129], and BCIN_16g01130, the bcpka1 catalytic subunit of the cAMP-dependent protein kinase (PKA), whose deletion affects the lesion development and leaves rot caused by the fungus [130]. Two other putative targets were BcnoxA (BCIN_05g00350), a component of the *B. cinerea* NADPH oxidase complex necessary for the colonization of host tissues [131], and the MAP triple kinase BcSte11 (BCIN_03g02630), whose deletion is known to cause defects in germination, delayed vegetative growth, reduced size of conidia, lack of sclerotia formation and loss of pathogenicity in *B. cinerea* [132]. Moreover, a *B. cinerea* homolog of *Ssams2* (BCIN_08g03180) was also among the putative targets, and this gene encodes a GATA transcription factor required for appressoria formation and chromosome segregation in *S. sclerotiorum* [115].

Several other genes encoding virulence factors were found to be upregulated in the pathogenic mycohosts during the interaction with the Δ*dcl2* mutant, even if they were not among the putative targets of milRNAs (Table 6). Among the *F. graminearum* genes upregulated during contact with Δ*dcl2* were the transcription factors MYT3, ERB1, GzHMG005, GzC2H066, GzZC230 and GzC2H059, whose disruption reduces the virulence of the pathogen [126,133– 136], as well as the phospholipase PLC1, known for its involvement in hyphal growth, conidiation, deoxynivalenol production and virulence [137]. Regarding *B. cinerea*, among the genes upregulated during contact with the Δ*dcl2* mutant we find nop53 and noxR, crucial for fungal development and virulence through the regulation of reactive oxygen species [138, 139], frq1, involved in circadian regulation of fungal virulence [140], and vel1, whose deletion affects virulence and light-dependent differentiation [141]. Moreover, among the upregulated genes there was also a homolog (BCIN_14g03930) of the *S. sclerotiorum* transcription factor *SsNsd1*, necessary for pathogenicity and appressorium formation [142]. Furthermore, upon contact with the Δ*dcl2* mutant *B. cinerea* upregulated several genes encoding proteins involved in chitin and cell wall synthesis, such as Bccrh1, BcchsIV, BcchsV, BcchsVI and BcchsVII [143–146]. The upregulation of BcCHSVI and BcCHSVII is of particular interest because these proteins have a role in plant infection [146].

Genes encoding two virulence factors of *F. graminearum* (TRI5 and TRI6) and one of *B. cinerea* (MFSG) were downregulated during interaction with the Δ*dcl2* mutant. The gene *mfsG* is involved in *B. cinerea* virulence by providing tolerance to glucosinolate-breakdown products [147], but the *C. rosea* Δ*dcl2* mutant shows downregulation in several putative secondary metabolite clusters compared to the WT. Therefore, it is possible that the expression of *mfsG* is reduced during contact with the mutant because the lack of production of harmful compounds makes it unnecessary for the mycohost to express resistance genes. TRI5 and TRI6 are involved in the synthesis of trichothecenes [148, 149], and other genes involved in the biosynthesis of these mycotoxins are similarly downregulated during contact with the Δ*dcl2* mutant, including the genes *TRI1*, *TRI3*, *TRI4*, *TRI9* and *TRI14* [150]. This is surprising because *F. graminearum* overexpresses the transcription factor gene *ART1* during contact with the Δ*dcl2* mutant, and this transcription factor is known to be a positive regulator of trichothecene biosynthesis [128]. The reduced ability of the Δ*dcl2* mutant to control *F. graminearum* may make it unnecessary for the mycohost to produce DON in high quantities, despite *ART1* overexpression. Interestingly, among the most relevant genes proven to be DON-responsive in *C. rosea* in a previous study [151], only one out of 16 was found to be less expressed in the Δ*dcl2* mutant than in the WT during interaction with *F. graminearum*: a homolog of glucose repressible protein GRG1 (CRV2G00000966). Given the reduced expression of DON-biosynthesis genes by *F. graminearum*, the downregulation of a higher number of DON-responsive genes was expected.

Another important mycotoxin produced by *F. graminearum* is zearalenone, and the zearalenone hydrolase gene *zhd101* (CRV2G00011056) was found to be downregulated by the Δ*dcl2* mutant. The deletion of this gene undermines *C. rosea* mycoparasitic action against *F. graminearum* [152] and its downregulation is, therefore, a possible reason for the impaired biocontrol action of the Δ*dcl2* mutant. Another zearalenone-responsive gene, encoding a putative bacteriorhodopsin-like protein [151], is also downregulated in the Δ*dcl2* mutant, but it’s role in the *C. rosea-F. graminearum* interaction is still unknown.

Interestingly, *F. graminearum* showed altered production of red pigment at the point of contact with the Δ*dcl2* mutant, which could plausibly be due to downregulation of genes belonging to the gene clusters of carotenoid and fusarielin [153, 154] (Table 6). However, the gene cluster of aurofusarins, known for their red colorations, was not differentially expressed during the interaction with Δ*dcl2*. It is also necessary to mention that *dcl2* deletion can impair the biocontrol activity of *C. rosea* through reducing its ability to induce plant defense responses. For example, the Δ*dcl2* mutant showed a reduced capacity of producing harzianolide D, which has been proven to be a systemic resistance elicitor on tomatoes [155].

## 5. Conclusions

DCL-dependent RNA silencing plays a determinant role in the regulation of many biological processes. In the present study, the role of DCL-like enzymes was investigated for the first time in the antagonistic action of the fungus *C. rosea*. Our result show that DCL2-mediated RNAi plays a central role in regulating endogenous cellular processes involved in growth, secondary metabolite production and antagonism towards the mycohosts, while function of DCL1 is redundant except for conidia production. The observed phenotypic effect in Δ*dcl2* strains is due to the diminished production of antifungal metabolites in the mutant, as well as to downregulation of genes known to be involved in mycohost response and resistance to secondary metabolites. Identification of 11 milRNAs, which were downregulated in Δ*dcl2* strain, and their putative endogenous gene targets including transcription factors and chromatin remodeling proteins indicates DCL-dependent regulation of *C. rosea* antagonistic interactions. Furthermore, we identified cross-species gene targets in the mycohosts *B. cinerea* and *F. graminearum* previously characterized for their role in fungal virulence. This highlights a possible cross-species regulation activity of *C. rosea* milRNAs, posing the bases for future studies focusing on the role of DCL-dependent RNA silencing in interspecific fungal interactions.

## 6. Acknowledgements

This work was financially supported by the Department of Forest Mycology and Plant Pathology; Swedish Research Council for Environment, Agricultural Sciences and Spatial Planning (FORMAS; grant number 2018-01420), and Carl Tryggers Stiftelse för Vetenskaplig Forskning (CTS 19: 82). MK acknowledges SLU Centre for Biological Control (CBC) at the Swedish University of Agricultural Sciences. RV is supported by FORMAS (2019-01316), Carl Tryggers Stiftelse för Vetenskaplig Forskning (CTS 20: 464) and The Crafoord foundation (20200818).

## 10. Supplementary information

**Figure S1:** Generation and validation of gene deletion and complementation strains.

**Figure S2:** Setup for dual culture interactions.

**Figure S3:** Diagrammatic representation of protein domains predicted by Simple Modular Architecture Research Tool (SMART) and Conserved Domain Database (CDD) within Dicer-like (A), argonaut (B) and RNA-dependent RNA polymerase (C) proteins.

**Figure S4:** phylogenetic trees presenting the evolutionary relationship among dicer like proteins (**A**), argonauts (**B**) and RNA dependent RNA polymerases (**C**) of several fungal plant pathogens. The trees were generated with iqtree v.1.6.12 and visualized with figtree v.1.4.4.

**Figure S5:** Phenotypic characterization of *C*. *rosea* strains.

**Figure S6:** Heatmaps showing all compounds significantly underproduced in the Δ*dcl1*/Δ*dcl2* strains compared to WT and significantly overproduced in the Δ*dcl1*+/Δ*dcl2+* strains compared to the Δ*dcl1*/Δ*dcl2* strains. (**A**) WT/Δ*dcl1*/Δ*dcl1+*. (**B**) WT/Δ*dcl2*/Δ*dcl2+*. Dark red: high concentration. White: low concentration.

**Figure S7:** Tentative identification of selected sorbicillin type compounds by UHPLC-MSMS. (**A**) Selected MSMS spectra. (**B-F**) Proposed formation of observed fragment ions for oxosorbicillinol, sorbicillinol, epoxysorbicillinol, sorbicillin and bisvertinolone, respectively.

**Table S1:** Protein ID of the putative argonauts, dicer like proteins and RNA dependent RNA polymerases proteins used in phylogenetic analyses.

**Table S2:** List of primers used in this study.

**Table S3:** Characteristics of the putative argonauts dicer like proteins and RNA dependent RNA polymerases proteins in *Clonostachys rosea*.

**Table S4:** Secondary metabolite analysis of *C*. *rosea* WT, Δ*dcl1* and Δ*dcl2*, and Δ*dcl1*+ and Δ*dcl2**+*** strains by UHPLC-MS and -MSMS.

**Table S5:** Results of STAR and featureCounts based mapping of mRNA reads. The reads were mapped to the concatenated genomes of *C. rosea* and the mycohot interacting with it, which was either *B. cinerea* or *F. graminearum*, depending on the sample.

**Table S6:** Differentially expressed *C. rosea* genes during the interspecies interactions.

**Table S7:** Expression and annotation of MFS transporters, secondary metabolite gene clusters, glycoside hydrolases, ABC transporters and transcription factors differentially expressed in this study.

**Table S8:** C. rosea genes associated with gene silencing machinery and chromatin modification.

**Table S9:** Results of STAR and featureCounts based mapping of sRNA reads. The reads were mapped to the concatenated genomes of *C. rosea* and the mycohot interacting with it, which was either *B. cinerea* or *F. graminearum*, depending on the sample.

**Table S10:** Sequence, location, and expression values of milRNAs detected in the study.

**Table S11:** Annotation and expression level of mycohot genes overexpressed during contact between *C. rosea* Δ*dcl2* mutant and the pathogens, putatively targeted by *C. rosea* milRNAs underexpressed in the same interaction.

**Table S12:** Differentially expressed *B*. *cinerea* and *F*. *graminearum* genes during the interaction with *C*. *rosea* WT, Δ*dcl1* and Δ*dcl2* strains.

## References

1. Ghildiyal M, Zamore PD. Small silencing RNAs: An expanding universe. Nat Rev Genet. 2009;10: 94–108. doi:10.1038/nrg2504

2. Hannon GJ. RNA interference. Nature. 2002;418: 244–251. doi:10.1038/418244a

3. Huang CY, Wang H, Hu P, Hamby R, Jin H. Small RNAs – Big Players in Plant-Microbe Interactions. Cell Host Microbe. 2019;26: 173–182. doi:10.1016/j.chom.2019.07.021

4. Malone CD, Hannon GJ. Small RNAs as Guardians of the Genome. Cell. 2009;136: 656–668. doi:10.1016/j.cell.2009.01.045

5. Van Wolfswinkel JC, Ketting RF. The role of small non-coding RNAs in genome stability and chromatin organization. J Cell Sci. 2010;123: 1825–1839. doi:10.1242/jcs.061713

6. Lee HC, Li L, Gu W, Xue Z, Crosthwaite SK, Pertsemlidis A, et al. Diverse Pathways Generate MicroRNA-like RNAs and Dicer-Independent Small Interfering RNAs in Fungi. Mol Cell. 2010;38: 803–814. doi:10.1016/j.molcel.2010.04.005

7. Nicolás FE, Ruiz-Vázquez RM. Functional diversity of RNAi-associated sRNAs in fungi. Int J Mol Sci. 2013;14: 15348–60. doi:10.3390/ijms140815348

8. Torres-Martínez S, Ruiz-Vázquez RM. The RNAi Universe in Fungi: A Varied Landscape of Small RNAs and Biological Functions. Annu Rev Microbiol. 2017;71: 371–391. doi:10.1146/annurev-micro-090816-093352

9. Romano N, Macino G. Quelling: transient inactivation of gene expression in Neurospora crassa by transformation with homologous sequences. Mol Microbiol. 1992;6: 3343–3353. doi:10.1111/j.1365-2958.1992.tb02202.x

10. Aramayo R, Metzenberg RL. Meiotic transvection in fungi. Cell. 1996;86: 103–113. doi:10.1016/S0092-8674(00)80081-1

11. Shiu PKT, Raju NB, Zickler D, Metzenberg RL. Meiotic silencing by unpaired DNA. Cell. 2001;107: 905–916. doi:10.1016/S0092-8674(01)00609-2

12. Hammond TM, Keller NP. RNA silencing in Aspergillus nidulans is independent of RNA-dependent RNA polymerases. Genetics. 2005;169: 607–617. doi:10.1534/genetics.104.035964

13. Janbon G, Maeng S, Yang DH, Ko YJ, Jung KW, Moyrand F, et al. Characterizing the role of RNA silencing components in Cryptococcus neoformans. Fungal Genet Biol. 2010;47: 1070–1080. doi:10.1016/j.fgb.2010.10.005

14. Wang X, Wang P, Sun S, Darwiche S, Idnurm A, Heitman J. Transgene Induced Co-Suppression during Vegetative Growth in Cryptococcus neoformans. PLoS Genet. 2012;8: e1002885. doi:10.1371/journal.pgen.1002885

15. Son H, Min K, Lee J, Raju NB, Lee YW. Meiotic silencing in the homothallic fungus Gibberella zeae. Fungal Biol. 2011;115: 1290–1302. doi:10.1016/j.funbio.2011.09.006

16. Kadotani N, Nakayashiki H, Tosa Y, Mayama S. RNA silencing in the phytopathogenic fungus Magnaporthe oryzae. Mol Plant-Microbe Interact. 2003;16: 769–776. doi:10.1094/MPMI.2003.16.9.769

17. Murata T, Kadotani N, Yamaguchi M, Tosa Y, Mayama S, Nakayashiki H. siRNA-dependent and-independent post-transcriptional cosuppression of the LTR-retrotransposon MAGGY in the phytopathogenic fungus Magnaporthe oryzae. Nucleic Acids Res. 2007;35: 5987–5994.

18. De Haro JP, Calo S, Cervantes M, Nicolás FE, Torres-Martinez S, Ruiz-Vázquez RM. A single dicer gene is required for efficient gene silencing associated with two classes of small antisense RNAs in mucor circinelloides. Eukaryot Cell. 2009;8: 1486–1497. doi:10.1128/EC.00191-09

19. Nicolas FE, Moxon S, de Haro JP, Calo S, Grigoriev I V., Torres-MartÍnez S, et al. Endogenous short RNAs generated by Dicer 2 and RNA-dependent RNA polymerase 1 regulate mRNAs in the basal fungus Mucor circinelloides. Nucleic Acids Res. 2010;38: 5535–5541. doi:10.1093/nar/gkq301

20. Carreras-Villaseñor N, Esquivel-Naranjo EU, Villalobos-Escobedo JM, Abreu-Goodger C, Herrera-Estrella A. The RNAi machinery regulates growth and development in the filamentous fungus Trichoderma atroviride. Mol Microbiol. 2013. doi:10.1111/mmi.12261

21. Laurie JD, Linning R, Bakkeren G. Hallmarks of RNA silencing are found in the smut fungus Ustilago hordei but not in its close relative Ustilago maydis. Curr Genet. 2008;53: 49–58. doi:10.1007/s00294-007-0165-7

22. Drinnenberg IA, Weinberg DE, Xie KT, Mower JP, Wolfe KH, Fink GR, et al. RNAi in budding yeast. Science (80-).1 2009;326: 544–550. doi:10.1126/science.1176945

23. Dang Y, Yang Q, Xue Z, Liu Y. RNA interference in fungi: Pathways, functions, and applications. Eukaryot Cell. 2011;10: 1148–1155. doi:10.1128/EC.05109-11

24. Bai Y, Lan F, Yang W, Zhang F, Yang K, Li Z, et al. SRNA profiling in Aspergillus flavus reveals differentially expressed miRNA-like RNAs response to water activity and temperature. Fungal Genet Biol. 2015;81: 113–119. doi:10.1016/j.fgb.2015.03.004

25. Kang K, Zhong J, Jiang L, Liu G, Gou CY, Wu Q, et al. Identification of microRNA-Like RNAs in the Filamentous Fungus Trichoderma reesei by Solexa Sequencing. PLoS One. 2013;8: e76288. doi:10.1371/journal.pone.0076288

26. Zhou J, Fu Y, Xie J, Li B, Jiang D, Li G, et al. Identification of microRNA-like RNAs in a plant pathogenic fungus Sclerotinia sclerotiorum by high-throughput sequencing. Mol Genet Genomics. 2012;287: 275–282. doi:10.1007/s00438-012-0678-8

27. Lau SKP, Chow WN, Wong AYP, Yeung JMY, Bao J, Zhang N, et al. Identification of MicroRNA-Like RNAs in Mycelial and Yeast Phases of the Thermal Dimorphic Fungus Penicillium marneffei. PLoS Negl Trop Dis. 2013;7: e2398. doi:10.1371/journal.pntd.0002398

28. Lin YL, Ma LT, Lee YR, Lin SS, Wang SY, Chang TT, et al. MicroRNA-like small RNAs prediction in the development of antrodia cinnamomea. PLoS One. 2015;10: e0123245. doi:10.1371/journal.pone.0123245

29. Lin R, He L, He J, Qin P, Wang Y, Deng Q, et al. Comprehensive analysis of microRNA-seq and target mRNAs of rice sheath blight pathogen provides new insights into pathogenic regulatory mechanisms. DNA Res. 2016;23: 415–425. doi:10.1093/dnares/dsw024

30. Liu T, Hu J, Zuo Y, Jin Y, Hou J. Identification of microRNA-like RNAs from Curvularia lunata associated with maize leaf spot by bioinformation analysis and deep sequencing. Mol Genet Genomics. 2016;291: 587–596. doi:10.1007/s00438-015-1128- 1

31. Mueth NA, Ramachandran SR, Hulbert SH. Small RNAs from the wheat stripe rust fungus (Puccinia striiformis f.sp. tritici). BMC Genomics. 2015;16:1–16. doi:10.1186/s12864-015-1895-4

32. Nunes CC, Gowda M, Sailsbery J, Xue M, Chen F, Brown DE, et al. Diverse and tissue-enriched small RNAs in the plant pathogenic fungus, Magnaporthe oryzae. BMC Genomics. 2011;12: 1–20. doi:10.1186/1471-2164-12-288

33. Raman V, Simon SA, Romag A, Demirci F, Mathioni SM, Zhai J, et al. Physiological stressors and invasive plant infections alter the small RNA transcriptome of the rice blast fungus, Magnaporthe oryzae. BMC Genomics. 2013;14: 1–18. doi:10.1186/1471-2164-14-326

34. Zhou Q, Wang Z, Zhang J, Meng H, Huang B. Genome-wide identification and profiling of microRNA-like RNAs from Metarhizium anisopliae during development. Fungal Biol. 2012;116: 1156–1162. doi:10.1016/j.funbio.2012.09.001

35. Weiberg A, Wang M, Lin FM, Zhao H, Zhang Z, Kaloshian I, et al. Fungal small RNAs suppress plant immunity by hijacking host RNA interference pathways. Science (80-). 2013;342: 118–123. doi:10.1126/science.1239705

36. Weiberg A, Jin H. Small RNAs-the secret agents in the plant-pathogen interactions. Curr Opin Plant Biol. 2015;26. doi:10.1016/j.pbi.2015.05.033

37. Chaloner T, van Kan JAL, Grant-Downton RT. RNA ‘Information Warfare’ in Pathogenic and Mutualistic Interactions. Trends Plant Sci. 2016;21: 738–748. doi:10.1016/j.tplants.2016.05.008

38. Wang M, Weiberg A, Lin FM, Thomma BPHJ, Huang H Da, Jin H. Bidirectional cross-kingdom RNAi and fungal uptake of external RNAs confer plant protection. Nat Plants. 2016;2: 1–10. doi:10.1038/nplants.2016.151

39. Zhang T, Zhao YL, Zhao JH, Wang S, Jin Y, Chen ZQ, et al. Cotton plants export microRNAs to inhibit virulence gene expression in a fungal pathogen. Nat Plants. 2016;2: 1–6. doi:10.1038/nplants.2016.153

40. Cui C, Wang Y, Liu J, Zhao J, Sun P, Wang S. A fungal pathogen deploys a small silencing RNA that attenuates mosquito immunity and facilitates infection. Nat Commun. 2019;10: 1–10. doi:10.1038/s41467-019-12323-1

41. Karlsson M, Durling MB, Choi J, Kosawang C, Lackner G, Tzelepis GD, et al. Insights on the evolution of mycoparasitism from the genome of clonostachys rosea. Genome Biol Evol. 2015. doi:10.1093/gbe/evu292

42. Iqbal M, Dubey M, Mcewan K, Menzel U, Franko MA, Viketoft M, et al. Evaluation of clonostachys rosea for control of plant-parasitic nematodes in soil and in roots of carrot and wheat. Phytopathology. 2018;108: 52–59. doi:10.1094/PHYTO-03-17-0091-R

43. Dubey M, Vélëz H, Broberg M, Jensen DF, Karlsson M. LysM Proteins Regulate Fungal Development and Contribute to Hyphal Protection and Biocontrol Traits in Clonostachys rosea. Front Microbiol. 2020;11: 679. doi:10.3389/fmicb.2020.00679

44. Tzelepis G, Dubey M, Jensen DF, Karlsson M. Identifying glycoside hydrolase family 18 genes in the mycoparasitic fungal species clonostachys rosea. Microbiol (United Kingdom). 2015. doi:10.1099/mic.0.000096

45. Dubey MK, Jensen DF, Karlsson M. Hydrophobins are required for conidial hydrophobicity and plant root colonization in the fungal biocontrol agent Clonostachys rosea. BMC Microbiol. 2014;14: 1–14. doi:10.1186/1471-2180-14-18

46. Sun Z Bin, Sun MH, Li SD. Identification of mycoparasitism-related genes in Clonostachys rosea 67-1 active against Sclerotinia sclerotiorum. Sci Rep. 2015. doi:10.1038/srep18169

47. Demissie ZA, Foote SJ, Tan Y, Loewen MC. Profiling of the transcriptomic responses of Clonostachys rosea Upon treatment with Fusarium graminearum Secretome. Front Microbiol. 2018;9: 1061. doi:10.3389/fmicb.2018.01061

48. Demissie ZA, Witte T, Robinson KA, Sproule A, Foote SJ, Johnston A, et al. Transcriptomic and Exometabolomic Profiling Reveals Antagonistic and Defensive Modes of Clonostachys rosea Action against Fusarium graminearum. Mol Plant-Microbe Interact. 2020;33: 842–858. doi:10.1094/MPMI-11-19-0310-R

49. Nygren K, Dubey M, Zapparata A, Iqbal M, Tzelepis GD, Durling MB, et al. The mycoparasitic fungus Clonostachys rosea responds with both common and specific gene expression during interspecific interactions with fungal prey. Evol Appl. 2018;11: 931–949. doi:10.1111/eva.12609

50. Fatema U, Broberg A, Jensen DF, Karlsson M, Dubey M. Functional analysis of polyketide synthase genes in the biocontrol fungus Clonostachys rosea. Sci Rep. 2018;8: 1–17. doi:10.1038/s41598-018-33391-1

51. Dubey M, Jensen D, Karlsson M. The ABC transporter ABCG29 is involved in H2O2 tolerance and biocontrol traits in the fungus clonostachys rosea. Mol Genet Genomics. 2016;291: 677–686. doi:10.1007/s00438-015-1139-y

52. Kamou NN, Dubey M, Tzelepis G, Menexes G, Papadakis EN, Karlsson M, et al. Investigating the compatibility of the biocontrol agent Clonostachys rosea IK726 with prodigiosin-producing Serratia rubidaea S55 and phenazine-producing Pseudomonas chlororaphis ToZa7. Arch Microbiol. 2016;198: 369–377. doi:10.1007/s00203-016-1198-4

53. Dubey MK, Jensen DF, Karlsson M. An ATP-binding cassette pleiotropic drug transporter protein is required for xenobiotic tolerance and antagonism in the fungal biocontrol agent Clonostachys rosea. Mol Plant-Microbe Interact. 2014;27: 725–732. doi:10.1094/MPMI-12-13-0365-R

54. Iqbal M, Broberg M, Haarith D, Broberg A, Bushley KE, Brandström Durling M, et al. Natural variation of root lesion nematode antagonism in the biocontrol fungus Clonostachys rosea and identification of biocontrol factors through genome-wide association mapping. Evol Appl. 2020;13: 2264. doi:10.1111/eva.13001

55. Broberg M, Dubey M, Sun MH, Ihrmark K, Schroers HJ, Li SD, et al. Out in the cold: Identification of genomic regions associated with cold tolerance in the biocontrol fungus clonostachys roseathrough genome-wide association mapping. Front Microbiol. 2018;9: 2844. doi:10.3389/fmicb.2018.02844

56. Letunic I, Doerks T, Bork P. SMART 6: Recent updates and new developments. Nucleic Acids Res. 2009;37: 229–232. doi:10.1093/nar/gkn808

57. Jones P, Binns D, Chang HY, Fraser M, Li W, McAnulla C, et al. InterProScan 5: Genome-scale protein function classification. Bioinformatics. 2014;30: 1236–1240. doi:10.1093/bioinformatics/btu031

58. Marchler-Bauer A, Lu S, Anderson JB, Chitsaz F, Derbyshire MK, DeWeese-Scott C, et al. CDD: A Conserved Domain Database for the functional annotation of proteins. Nucleic Acids Res. 2011;39: 225–229. doi:10.1093/nar/gkq1189

59. Bateman A, Martin MJ, O’Donovan C, Magrane M, Apweiler R, Alpi E, et al. UniProt: A hub for protein information. Nucleic Acids Res. 2015. doi:10.1093/nar/gku989

60. Benson DA, Cavanaugh M, Clark K, Karsch-Mizrachi I, Ostell J, Pruitt KD, et al. GenBank. Nucleic Acids Res. 2018. doi:10.1093/nar/gkx1094

61. Katoh K, Standley DM. MAFFT multiple sequence alignment software version 7: Improvements in performance and usability. Mol Biol Evol. 2013. doi:10.1093/molbev/mst010

62. Nguyen LT, Schmidt HA, Von Haeseler A, Minh BQ. IQ-TREE: A fast and effective stochastic algorithm for estimating maximum-likelihood phylogenies. Mol Biol Evol. 2015. doi:10.1093/molbev/msu300

63. Rambaut A. FigTree v. 1.4.4. http://tree.bio.ed.ac.uk/software/figtree/. 2018.

64. Dubey MK, Ubhayasekera W, Sandgren M, Funck Jensen D, Karlsson M. Disruption of the Eng18b ENGase gene in the fungal biocontrol agent Trichoderma atroviride affects growth, conidiation and antagonistic ability. PLoS One. 2012;7: e36152. doi:10.1371/journal.pone.0036152

65. Güldener U, Heck S, Fiedler T, Beinhauer J, Hegemann JH. A new efficient gene disruption cassette for repeated use in budding yeast. Nucleic Acids Res. 1996;24: 2519–2524.

66. Karimi M, De Meyer B, Hilson P. Modular cloning in plant cells. Trends Plant Sci. 2005;10: 103–105. doi:10.1016/j.tplants.2005.01.008

67. Utermark J, Karlovsky P. Genetic transformation of filamentous fungi by Agrobacterium tumefaciens. Protoc Exch. 2008;119: 631–640. doi:10.1038/nprot.2008.83

68. Dubey MK, Broberg A, Sooriyaarachchi S, Ubhayasekera W, Jensen DF, Karlsson M. The glyoxylate cycle is involved in pleotropic phenotypes, antagonism and induction of plant defence responses in the fungal biocontrol agent Trichoderma atroviride. Fungal Genet Biol. 2013;58: 33–41. doi:10.1016/j.fgb.2013.06.008

69. Knudsen IMB, Hockenhull J, Jensen DF. Biocontrol of seedling diseases of barley and wheat caused by Fusarium culmorum and Bipolaris sorokiniana: effects of selected fungal antagonists on growth and yield components. Plant Pathol. 1995;44: 467–477. doi:10.1111/j.1365-3059.1995.tb01669.x

70. Smith CA, Want EJ, O’Maille G, Abagyan R, Siuzdak G. XCMS: processing mass spectrometry data for metabolite profiling using nonlinear peak alignment, matching, and identification. Anal Chem. 2006;78: 779–787.

71. Tautenhahn R, Böttcher C, Neumann S. Highly sensitive feature detection for high resolution LC/MS. BMC Bioinformatics. 2008;9: 1–16.

72. Hosseini P, Tremblay A, Matthews BF, Alkharouf NW. An efficient annotation and gene-expression derivation tool for Illumina Solexa datasets. BMC Res Notes. 2010;3: 1–7.

73. Conesa A, Götz S, García-Gómez JM, Terol J, Talón M, Robles M. Blast2GO: A universal tool for annotation, visualization and analysis in functional genomics research. Bioinformatics. 2005;21: 3674–3676. doi:10.1093/bioinformatics/bti610

74. Blin K, Shaw S, Steinke K, Villebro R, Ziemert N, Lee SY, et al. AntiSMASH 5.0: Updates to the secondary metabolite genome mining pipeline. Nucleic Acids Res. 2019;47: 81–87. doi:10.1093/nar/gkz310

75. Zhang H, Yohe T, Huang L, Entwistle S, Wu P, Yang Z, et al. DbCAN2: A meta server for automated carbohydrate-active enzyme annotation. Nucleic Acids Res. 2018;46: 95–101. doi:10.1093/nar/gky418

76. Urban M, Cuzick A, Rutherford K, Irvine A, Pedro H, Pant R, et al. PHI-base: A new interface and further additions for the multi-species pathogen-host interactions database. Nucleic Acids Res. 2017;45: 604–610. doi:10.1093/nar/gkw1089

77. Bushnell B. BBTools: A Suite of Fast, Multithreaded Bioinformatics Tools Designed for Analysis of DNA and RNA Sequence Data. Jt Genome Inst Berkeley, CA, USA, 2018. 2019.

78. Dobin A, Davis CA, Schlesinger F, Drenkow J, Zaleski C, Jha S, et al. STAR: Ultrafast universal RNA-seq aligner. Bioinformatics. 2013;29: 15–21. doi:10.1093/bioinformatics/bts635

79. Liao Y, Smyth GK, Shi W. FeatureCounts: An efficient general purpose program for assigning sequence reads to genomic features. Bioinformatics. 2014;30: 923–930. doi:10.1093/bioinformatics/btt656

80. Love MI, Anders S, Huber W. Differential analysis of count data - the DESeq2 package. Genome Biol. 2014;15: 10–1186.

81. Kopylova E, Noé L, Touzet H. SortMeRNA: Fast and accurate filtering of ribosomal RNAs in metatranscriptomic data. Bioinformatics. 2012;28: 3211–3217. doi:10.1093/bioinformatics/bts611

82. Quast C, Pruesse E, Yilmaz P, Gerken J, Schweer T, Yarza P, et al. The SILVA ribosomal RNA gene database project: Improved data processing and web-based tools. Nucleic Acids Res. 2013;41: 590–596. doi:10.1093/nar/gks1219

83. Paschoal AR, Maracaja-Coutinho V, Setubal JC, Simões ZLP, Verjovski-Almeida S, Durham AM. Non-coding transcription characterization and annotation: A guide and web resource for non-coding RNA databases. RNA Biol. 2012;9: 274–282. doi:10.4161/rna.19352

84. Gremme G, Steinbiss S, Kurtz S. Genome tools: A comprehensive software library for efficient processing of structured genome annotations. IEEE/ACM Trans Comput Biol Bioinforma. 2013;10: 645–656. doi:10.1109/TCBB.2013.68

85. Mackowiak SD. Identification of novel and known unit 12.10 miRNAs in deep-sequencing data with miRDeep2. Curr Protoc Bioinforma. 2011;36: 12–10. doi:10.1002/0471250953.bi1210s36

86. Kozomara A, Birgaoanu M, Griffiths-Jones S. MiRBase: From microRNA sequences to function. Nucleic Acids Res. 2019;47: 155–162. doi:10.1093/nar/gky1141

87. The RNA Central Consortium. RNAcentral: a hub of information for non-coding RNA sequences. Nucleic Acids Res. 2019;47: D221–D229.

88. Chen R, Jiang N, Jiang Q, Sun X, Wang Y, Zhang H, et al. Exploring microRNA-like small RNAs in the filamentous fungus Fusarium oxysporum. PLoS One. 2014;9: e104956. doi:10.1371/journal.pone.0104956

89. Devers EA, Branscheid A, May P, Krajinski F. Stars and symbiosis: Microrna- and microrna*-mediated transcript cleavage involved in arbuscular mycorrhizal symbiosis. Plant Physiol. 2011;156: 1990–2010. doi:10.1104/pp.111.172627

90. Wang L, Xu X, Yang J, Chen L, Liu B, Liu T, et al. Integrated microRNA and mRNA analysis in the pathogenic filamentous fungus Trichophyton rubrum. BMC Genomics. 2018;19: 1–14. doi:10.1186/s12864-018-5316-3

91. Xia Z, Wang Z, Kav NNV, Ding C, Liang Y. Characterization of microRNA-like RNAs associated with sclerotial development in Sclerotinia sclerotiorum. Fungal Genet Biol. 2020;144: 103471. doi:10.1016/j.fgb.2020.103471

92. Rueda A, Barturen G, Lebrón R, Gómez-Martín C, Alganza Á, Oliver JL, et al. SRNAtoolbox: An integrated collection of small RNA research tools. Nucleic Acids Res. 2015;43: 473. doi:10.1093/nar/gkv555

93. Enright A, John B, Gaul U, Tuschl T, Sander C, Marks D. MicroRNA targets in Drosophila. Genome Biol. 2003;4: 1–27.

94. Kertesz M, Iovino N, Unnerstall U, Gaul U, Segal E. The role of site accessibility in microRNA target recognition. Nat Genet. 2007;39: 1278–1284.

95. Sturm M, Hackenberg M, Langenberger D, Frishman D. TargetSpy: a supervised machine learning approach for microRNA target prediction. BMC Bioinformatics. 2010;11: 1–17.

96. Wu H-J, Ma Y-K, Chen T, Wang M, Wang X-J. PsRobot: a web-based plant small RNA meta-analysis toolbox. Nucleic Acids Res. 2012;40: W22–W28.

97. Bonnet E, He Y, Billiau K, Van de Peer Y. TAPIR, a web server for the prediction of plant microRNA targets, including target mimics. Bioinformatics. 2010;26: 1566– 1568.

98. Amselem J, Cornut G, Choisne N, Alaux M, Alfama-Depauw F, Jamilloux V, et al. RepetDB: A unified resource for transposable element references. Mob DNA. 2019;10: 1–8. doi:10.1186/s13100-019-0150-y

99. Kodama H, Komamine A. RNAi and Plant Gene Function Analysis. Springer; 2011.

100. Livak KJ, Schmittgen TD. Analysis of relative gene expression data using real-time quantitative PCR and the 2− ΔΔCT method. methods. 2001;25: 402–408.

101. Meng J, Wang X, Xu D, Fu X, Zhang X, Lai D, et al. Sorbicillinoids from fungi and their bioactivities. Molecules. 2016;21: 715.

102. Brown DW, Lee SH, Kim LH, Ryu JG, Lee S, Seo Y, et al. Identification of a 12-gene fusaric acid biosynthetic gene cluster in Fusarium species through comparative and functional genomics. Mol Plant-Microbe Interact. 2015;28: 319–332. doi:10.1094/MPMI-09-14-0264-R

103. Crutcher FK, Liu J, Puckhaber LS, Stipanovic RD, Bell AA, Nichols RL. FUBT, a putative MFS transporter, promotes secretion of fusaric acid in the cotton pathogen Fusarium oxysporum f. Sp. vasinfectum. Microbiol (United Kingdom). 2015;161: 875–883. doi:10.1099/mic.0.000043

104. Reeves CD, Hu Z, Reid R, Kealey JT. Genes for the biosynthesis of the fungal polyketides hypothemycin from Hypomyces subiculosus and radicicol from Pochonia chlamydosporia. Appl Environ Microbiol. 2008;74: 5121–5129. doi:10.1128/AEM.00478-08

105. Yu J, Bhatnagar D, Cleveland TE. Completed sequence of aflatoxin pathway gene cluster in Aspergillus parasiticus. FEBS Lett. 2004;564: 126–130. doi:10.1016/S0014-5793(04)00327-8

106. Keller NP. Fungal secondary metabolism: regulation, function and drug discovery. Nat Rev Microbiol. 2019;17: 167–180. doi:10.1038/s41579-018-0121-1

107. Derntl C, Rassinger A, Srebotnik E, Mach RL, Mach-Aigner AR. Identification of the main regulator responsible for synthesis of the typical yellow pigment produced by Trichoderma reesei. Appl Environ Microbiol. 2016;82: 6247–6257. doi:10.1128/AEM.01408-16

108. Derntl C, Guzmán-Chávez F, Mello-de-Sousa TM, Busse H-J, Driessen AJM, Mach RL, et al. In vivo study of the sorbicillinoid gene cluster in Trichoderma reesei. Front Microbiol. 2017;8: 2037.

109. Boylan MT, Mirabito PM, Willett CE, Zimmerman CR, Timberlake WE. Isolation and physical characterization of three essential conidiation genes from Aspergillus nidulans. Mol Cell Biol. 1987;7: 3113–3118. doi:10.1128/mcb.7.9.3113

110. Adams TH, Timberlake WE. Developmental repression of growth and gene expression in Aspergillus. Proc Natl Acad Sci U S A. 1990;87: 5405–5409. doi:10.1073/pnas.87.14.5405

111. Hindley J, Phear G, Stein M, Beach D. Sucl+ encodes a predicted 13-kilodalton protein that is essential for cell viability and is directly involved in the division cycle of Schizosaccharomyces pombe. Mol Cell Biol. 1987;7: 504–511. doi:10.1128/mcb.7.1.504

112. Finkel JS, Xu W, Huang D, Hill EM, Desai J V., Woolford CA, et al. Portrait of Candida albicans adherence regulators. PLoS Pathog. 2012;8: e1002525. doi:10.1371/journal.ppat.1002525

113. Ridenour JB, Bluhm BH. The HAP complex in Fusarium verticillioides is a key regulator of growth, morphogenesis, secondary metabolism, and pathogenesis. Fungal Genet Biol. 2014;69: 52–64. doi:10.1016/j.fgb.2014.05.003

114. Son H, Park AR, Lim JY, Shin C, Lee YW. Genome-wide exonic small interference RNA-mediated gene silencing regulates sexual reproduction in the homothallic fungus Fusarium graminearum. PLoS Genet. 2017;13: e1006595. doi:10.1371/journal.pgen.1006595

115. Liu L, Wang Q, Zhang X, Liu J, Zhang Y, Pan H. Ssams2, a gene encoding GATA transcription factor, is required for appressoria formation and chromosome segregation in sclerotinia sclerotiorum. Front Microbiol. 2018;9: 3031. doi:10.3389/fmicb.2018.03031

116. Hirayama S, Sugiura R, Lu Y, Maeda T, Kawagishi K, Yokoyama M, et al. Zinc finger protein Prz1 regulates Ca2+ but not Cl-homeostasis in fission yeast. Identification of distinct branches of calcineurin signaling pathway in fission yeast. J Biol Chem. 2003;278: 18078–18084. doi:10.1074/jbc.M212900200

117. Ryan DP, Owen-Hughes T. Snf2-family proteins: Chromatin remodellers for any occasion. Curr Opin Chem Biol. 2011;15: 649–656. doi:10.1016/j.cbpa.2011.07.022

118. Baccile JA, Spraker JE, Le HH, Brandenburger E, Gomez C, Bok JW, et al. Plant-like biosynthesis of isoquinoline alkaloids in Aspergillus fumigatus. Nat Chem Biol. 2016;12: 419–424. doi:10.1038/nchembio.2061

119. Macheleidt J, Scherlach K, Neuwirth T, Schmidt-Heck W, Straßburger M, Spraker J, et al. Transcriptome analysis of cyclic AMP-dependent protein kinase A-regulated genes reveals the production of the novel natural compound fumipyrrole by Aspergillus fumigatus. Mol Microbiol. 2015;86: 148–162. doi:10.1111/mmi.12926

120. Zhai MM, Qi FM, Li J, Jiang CX, Hou Y, Shi YP, et al. Isolation of Secondary Metabolites from the Soil-Derived Fungus Clonostachys rosea YRS-06, a Biological Control Agent, and Evaluation of Antibacterial Activity. J Agric Food Chem. 2016;64: 2298–2306. doi:10.1021/acs.jafc.6b00556

121. Seidl V, Huemer B, Seiboth B, Kubicek CP. A complete survey of Trichoderma chitinases reveals three distinct subgroups of family 18 chitinases. FEBS J. 2005;272: 5923–5939. doi:10.1111/j.1742-4658.2005.04994.x

122. Karlsson M, Stenlid J. Comparative evolutionary histories of the fungal Chitinase gene family reveal non-random size expansions and contractions due to adaptive natural selection. Evol Bioinforma. 2008;4: EBO-S604. doi:10.4137/ebo.s604

123. Steindorff AS, Ramada MHS, Coelho ASG, Miller RNG, Pappas GJ, Ulhoa CJ, et al. Identification of mycoparasitism-related genes against the phytopathogen Sclerotinia sclerotiorum through transcriptome and expression profile analysis in Trichoderma harzianum. BMC Genomics. 2014;61: 134–140. doi:10.1186/1471-2164-15-204

124. Broberg M, Dubey M, Iqbal M, Gudmundssson M, Ihrmark K, Schroers HJ, et al. Comparative genomics highlights the importance of drug efflux transporters during evolution of mycoparasitism in Clonostachys subgenus Bionectria (Fungi, Ascomycota, Hypocreales). Evol Appl. 2020;14: 476. doi:10.1111/eva.13134

125. Ridenour JB, Smith JE, Bluhm BH. The HAP complex governs fumonisin biosynthesis and maize kernel pathogenesis in Fusarium verticillioides. J Food Prot. 2016;79: 1498–1507. doi:10.4315/0362-028X.JFP-15-596

126. Son H, Seo YS, Min K, Park AR, Lee J, Jin JM, et al. A phenome-based functional analysis of transcription factors in the cereal head blight fungus, fusarium graminearum. PLoS Pathog. 2011;7: e1002310. doi:10.1371/journal.ppat.1002310

127. Seong K, Hou Z, Tracy M, Kistler HC, Xu JR. Random insertional mutagenesis identifies genes associated with virulence in the wheat scab fungus Fusarium graminearum. Phytopathology. 2005;95: 744–750. doi:10.1094/PHYTO-95-0744

128. Oh M, Son H, Choi GJ, Lee C, Kim JC, Kim H, et al. Transcription factor ART1 mediates starch hydrolysis and mycotoxin production in Fusarium graminearum and F.verticillioides. Mol Plant Pathol. 2016. doi:10.1111/mpp.12328

129. Gourgues M, Brunet-Simon A, Lebrun MH, Levis C. The tetraspanin BcPIs1 is required for appressorium-mediated penetration of Botrytis cinerea into host plant leaves. Mol Microbiol. 2004;51: 619–629. doi:10.1046/j.1365-2958.2003.03866.x

130. Schumacher J, Kokkelink L, Huesmann C, Jimenez-Teja D, Collado IG, Barakat R, et al. The cAMP-dependent signaling pathway and its role in conidial germination, growth, and virulence of the gray mold Botrytis cinerea. Mol Plant-Microbe Interact. 2008;21: 1443–1459. doi:10.1094/MPMI-21-11-1443

131. Siegmund U, Marschall R, Tudzynski P. BcNoxD, a putative ER protein, is a new component of the NADPH oxidase complex in Botrytis cinerea. Mol Microbiol. 2015;95: 988–1005. doi:10.1111/mmi.12869

132. Schamber A, Leroch M, Diwo J, Mendgen K, Hahn M. The role of mitogen-activated protein (MAP) kinase signalling components and the Ste12 transcription factor in germination and pathogenicity of Botrytis cinerea. Mol Plant Pathol. 2010;11: 105– 119. doi:10.1111/j.1364-3703.2009.00579.x

133. Kim Y, Kim H, Son H, Choi GJ, Kim JC, Lee YW. MYT3, a Myb-like transcription factor, affects fungal development and pathogenicity of Fusarium graminearum. PLoS One. 2014;9: e94359. doi:10.1371/journal.pone.0094359

134. Jonkers W, Xayamongkhon H, Haas M, Olivain C, van der Does HC, Broz K, et al. EBR1 genomic expansion and its role in virulence of Fusarium species. Environ Microbiol. 2014;16: 1982–2003. doi:10.1111/1462-2920.12331

135. Zhao C, Waalwijk C, De Wit PJGM, Van Der Lee T, Tang D. EBR1, a novel Zn2Cys6 transcription factor, affects virulence and apical dominance of the hyphal tip in Fusarium graminearum. Mol Plant-Microbe Interact. 2011;24: 1407–1418. doi:10.1094/MPMI-06-11-0158

136. Dufresne M, Lee T van der, M’Barek S Ben, Xu X, Zhang X, Liu T, et al. Transposon-tagging identifies novel pathogenicity genes in Fusarium graminearum. Fungal Genet Biol. 2008;45: 1552–1561. doi:10.1016/j.fgb.2008.09.004

137. Zhu Q, Sun L, Lian J, Gao X, Zhao L, Ding M, et al. The phospholipase C (FgPLC1) is involved in regulation of development, pathogenicity, and stress responses in Fusarium graminearum. Fungal Genet Biol. 2016;97: 1–9. doi:10.1016/j.fgb.2016.10.004

138. Cao SN, Yuan Y, Qin YH, Zhang MZ, de Figueiredo P, Li GH, et al. The pre-rRNA processing factor Nop53 regulates fungal development and pathogenesis via mediating production of reactive oxygen species. Environ Microbiol. 2018;20: 1531–1549. doi:10.1111/1462-2920.14082

139. Li H, Tian S, Qin G. NADPH oxidase ls crucial for the cellular redox homeostasis in fungal pathogen botrytis cinerea. Mol Plant-Microbe Interact. 2019;32: 1508–1516. doi:10.1094/MPMI-05-19-0124-R

140. Hevia MA, Canessa P, Müller-Esparza H, Larrondo LF. A circadian oscillator in the fungus Botrytis cinerea regulates virulence when infecting Arabidopsis thaliana. Proc Natl Acad Sci U S A. 2015;112: 8744–8749. doi:10.1073/pnas.1508432112

141. Schumacher J, Pradier JM, Simon A, Traeger S, Moraga J, Collado IG, et al. Natural Variation in the VELVET Gene bcvel1 Affects Virulence and Light-Dependent Differentiation in Botrytis cinerea. PLoS One. 2012;7: e47840. doi:10.1371/journal.pone.0047840

142. Li J, Mu W, Veluchamy S, Liu Y, Zhang Y, Pan H, et al. The GATA-type IVb zinc-finger transcription factor SsNsd1 regulates asexual–sexual development and appressoria formation in Sclerotinia sclerotiorum. Mol Plant Pathol. 2018;19: 1679– 1689. doi:10.1111/mpp.12651

143. Bi K, Scalschi L, Jaiswal GN, Fried R, Zhu W, Masrati G, et al. The Botrytis cinerea Crh transglycosylae is a cytoplasmic effector triggering plant cell death and defense response. bioRxiv. 2020.

144. Cui Z, Ding Z, Yang X, Wang K, Zhu T. Gene disruption and characterization of a class V chitin synthase in Botrytis cinerea. Can J Microbiol. 2009;55: 1267–1274. doi:10.1139/W09-076

145. Cui Z, Wang Y, Lei N, Wang K, Zhu T. Botrytis cinerea chitin synthase BcChsVI is required for normal growth and pathogenicity. Curr Genet. 2013;59: 119–128. doi:10.1007/s00294-013-0393-y

146. Morcx S, Kunz C, Choquer M, Assie S, Blondet E, Simond-Côte E, et al. Disruption of Bcchs4, Bcchs6 or Bcchs7 chitin synthase genes in Botrytis cinerea and the essential role of class VI chitin synthase (Bcchs6). Fungal Genet Biol. 2013;52: 1–8. doi:10.1016/j.fgb.2012.11.011

147. Vela-Corcía D, Aditya Srivastava D, Dafa-Berger A, Rotem N, Barda O, Levy M. MFS transporter from Botrytis cinerea provides tolerance to glucosinolate-breakdown products and is required for pathogenicity. Nat Commun. 2019;10: 1–11. doi:10.1038/s41467-019-10860-3

148. McDonald T, Brown D, Keller NP, Hammond TM. RNA silencing of mycotoxin production in Aspergillus and Fusarium species. Mol Plant-Microbe Interact. 2005;18: 539–545. doi:10.1094/MPMI-18-0539

149. Bönnighausen J, Schauer N, Schäfer W, Bormann J. Metabolic profiling of wheat rachis node infection by Fusarium graminearum – decoding deoxynivalenol-dependent susceptibility. New Phytol. 2019;221: 459–469. doi:10.1111/nph.15377

150. Alexander NJ, Proctor RH, McCormick SP. Genes, gene clusters, and biosynthesis of trichothecenes and fumonisins in Fusarium. Toxin Rev. 2009;28: 198–215. doi:10.1080/15569540903092142

151. Kosawang C, Karlsson M, Jensen DF, Dilokpimol A, Collinge DB. Transcriptomic profiling to identify genes involved in Fusarium mycotoxin Deoxynivalenol and Zearalenone tolerance in the mycoparasitic fungus Clonostachys rosea. BMC Genomics. 2014;15: 1–11. doi:10.1186/1471-2164-15-55

152. Kosawang C, Karlsson M, Vélëz H, Rasmussen PH, Collinge DB, Jensen B, et al. Zearalenone detoxification by zearalenone hydrolase is important for the antagonistic ability of Clonostachys rosea against mycotoxigenic Fusarium graminearum. Fungal Biol. 2014;118: 364–373. doi:10.1016/j.funbio.2014.01.005

153. Jin JM, Lee J, Lee YW. Characterization of carotenoid biosynthetic genes in the ascomycete Gibberella zeae. FEMS Microbiol Lett. 2010;302: 197–202. doi:10.1111/j.1574-6968.2009.01854.x

154. Sørensen JL, Hansen FT, Sondergaard TE, Staerk D, Lee TV, Wimmer R, et al. Production of novel fusarielins by ectopic activation of the polyketide synthase 9 cluster in Fusarium graminearum. Environ Microbiol. 2012;14: 1159–1170. doi:10.1111/j.1462-2920.2011.02696.x

155. Cai F, Yu G, Wang P, Wei Z, Fu L, Shen Q, et al. Harzianolide, a novel plant growth regulator and systemic resistance elicitor from Trichoderma harzianum. Plant Physiol Biochem. 2013;73: 106–113.

